# Categorical judgments do not modify sensory information in working memory

**DOI:** 10.1101/2020.06.15.152223

**Authors:** Long Luu, Alan A. Stocker

## Abstract

Categorical judgments can systematically bias the perceptual interpretation of stimulus features. However, it remained unclear whether categorical judgments directly modify working memory representations or, alternatively, generate these biases via an inference process down-stream from working memory. To address this question we ran two novel psychophysical experiments in which human subjects had to revert their categorical judgments about a stimulus feature, if incorrect based on feedback, before providing an estimate of the feature. If categorical judgments indeed directly altered sensory representations in working memory, subjects’ estimates should reflect some aspects of their initial (incorrect) categorical judgment in those trials.

We found no traces of the initial categorical judgment. Rather, subjects seem to be able to flexibly switch their categorical judgment if needed and use the correct corresponding categorical prior to properly perform feature inference. A cross-validated model comparison also revealed that feedback may lead to selective memory recall such that only memory samples that are consistent with the categorical judgment are accepted for the inference process. Our results suggest that categorical judgments do not modify sensory information in working memory but rather act as top-down expectation in the subsequent sensory recall and inference process down-stream from working memory.

## Introduction

Human visual perception is biased by the statistical regularities in the sensory input. Many studies have demonstrated that these biases can be well accounted for by assuming that perception is an inference process that optimally trades-off sensory uncertainty with objective prior expectations reflecting these statistical regularities ^1,2,3,4,5,6^.

However, perception is not only biased by objective, statistical prior expectations but also by an observer’s subjective categorical assessment of a visual feature. For example, in a task sequence where subjects first had to make a categorical judgment about a stimulus feature before providing an estimate of the feature value (motion direction ^7,8,9^; orientation ^10,11^; or numerosity ^9,12^), the resulting estimates showed systematic biases in favor of their preceding category choices. Similar bias patterns have also been found in sequential tasks that did not explicitly involve categorical judgments ^13,14,8,15,16^ or estimates ^17,18^, suggesting that these biases are a wide-spread and general phenomenon in hierarchical perceptual inference ^19^. A general theory proposes that these bias patterns emerge because inference is intrinsically a top-down process where low-level feature estimates are conditioned on the observer’s preceding, (explicit or implicit) high-level categorical judgment ^20,21,15^. Embodying this theory, a “self-consistent” Bayesian observer model ^20^ has performed remarkably well in quantitatively capturing the rich and diverse perceptual behavior of human subjects in these tasks ^10^. The model assumes that a categorical judgment acts as a subjective prior that conditions the inference process of the stimulus feature. The motivation for humans to employ such a conditioned inference strategy is unclear, in particular because it is sub-optimal with regard to estimation accuracy. However, common to all tasks described above is that they are sequential processes that operate over time and require a working memory representation of the stimulus. Top-down conditioning may help to restore working memory that is corrupted by noise ^10,15^ although a detailed analysis revealed that this does not necessarily lead to improved accuracy ^19^. One established benefit of conditioning is that the observer’s interpretation of the stimulus remains self-consistent across all levels of representation (e.g., from category to feature level) despite corruption by memory or late noise ^10^. Conditioning may thus reflect a general inference strategy of the mind to avoid dissonant interpretations of the world at any moment in time ^22,23^.

While visual working memory seems a fundamental component of the perceptual process, its role in shaping the characteristic perceptual behavior in above tasks has not been well studied. With the current work, we aimed at identifying how and at what level a categorical judgment interacts with working memory representations in a way that leads to the observed post-decision biases. Specifically, we tested whether a categorical judgment directly modifies sensory representations in working memory, or not. We used an experimental design similar to our previous study ^10^: After presentation of an orientation stimulus, subjects first made a categorical judgment about the overall orientation and then subsequently provided an estimate of the orientation. In contrast to our previous work, however, subjects were given valid feedback about their categorical judgment in each trial and were instructed to use it for the estimation task. If categorical judgments modified working memory representation, we would expect subjects’ estimates to reflect these modifications in trials in which they had to change their categorical judgment because their initial assessment was incorrect. Our experimental results did not show any evidence that a categorical judgment updates and modifies working memory representations. Rather, subjects seemed to be able to flexibly change their categorical judgment if necessary (due to feedback) and combine it with stimulus information stored in working memory as predicted by the self-consistent observer model ^10^. This suggests that categorical and feature information have rather independent neural representations in working memory, which can be flexibly re-combined in the perceptual inference process if necessary.

## Results

Figure 1a outlines the two complementary hypotheses we tested. Based on the sensory information provided by a stimulus an observer first performs an implicit or explicit high-level categorical judgment about the stimulus. Stimulus information is stored in working memory and then used by the observer to make an estimate of the stimulus feature that, however, is biased by the preceding categorical decision. We previously demonstrated that a “self-consistent” Bayesian observer model can accurately account for human subjects’ behavior both in the decision (if explicit) and the estimation task ^21,10^. In this model, the observer’s categorical judgment acts as a prior which conditions feature inference to be consistent under this prior (see Methods for more details). However, what remained unclear was whether stimulus information (i.e. the likelihood function) and the categorical judgment (i.e. the prior) in each trial are separately represented and maintained and then independently combined down-stream of their working memory representation (*Hypothesis 2*), or whether the working memory representation is already reflecting a non-separable combination of the two (e.g. the posterior probability; *Hypothesis 1*).

**Figure 1:**
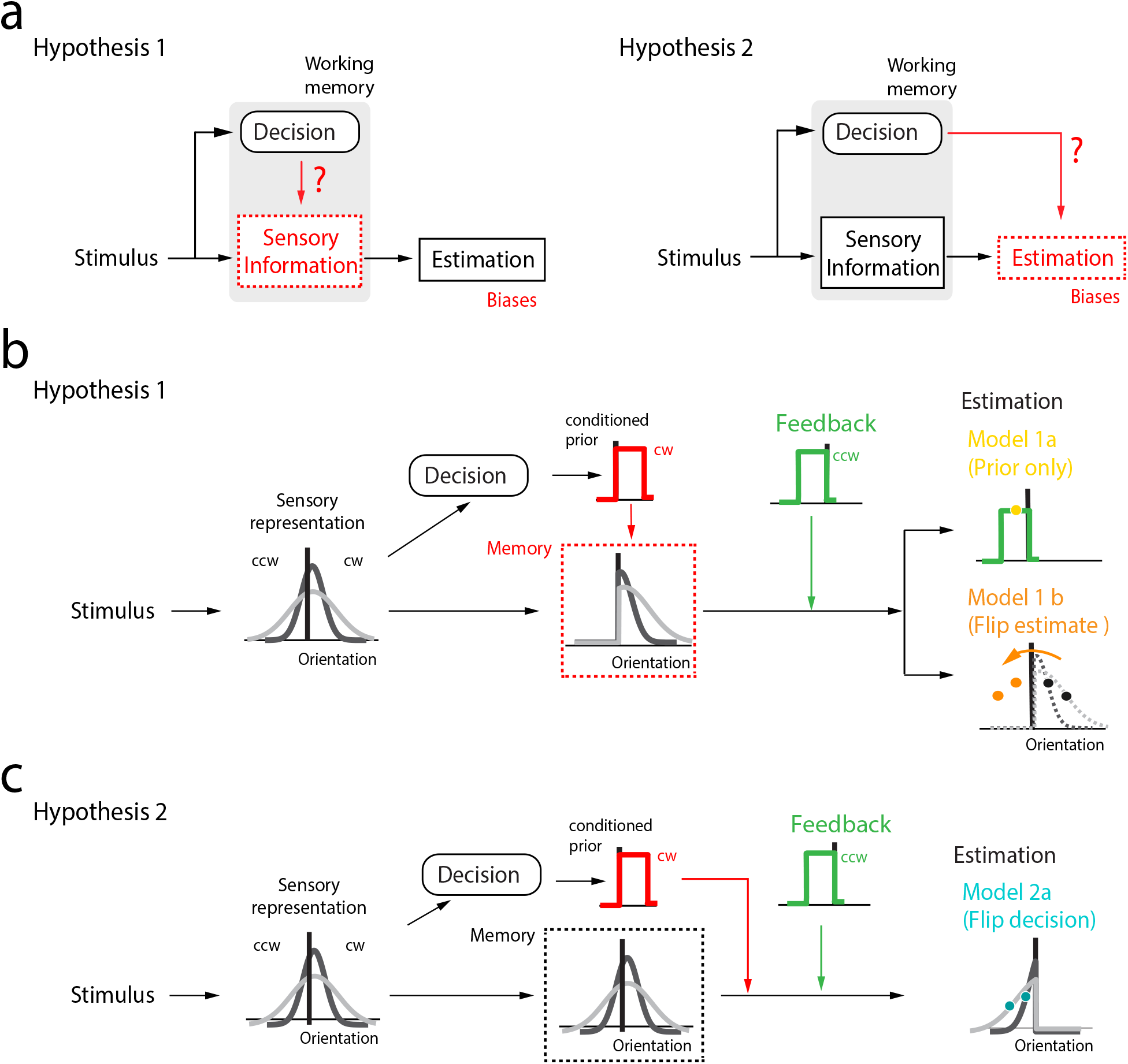
Potential mechanisms underlying choice-induced estimation biases. (**a**) Are choice-induced estimation biases the result of a modified sensory representation in working memory (Hypothesis 1) or rather of a computational inference process downstream (Hypothesis 2)? We run psychophysical experiments in which subjects had to reverse their categorical decision if incorrect according to feedback. (**b**) Hypothesis 1: After making a categorical judgment, working memory is updated to only reflect the sensory information consistent with the decision (i.e. the posterior). Upon receiving feedback indicating that the decision was incorrect, the conditioned prior is updated. Model 1a (Prior only) only relies on the updated prior for the estimate as the posterior does not provide any information on the correct side of the reference. Model 1b (Flip estimate) assumes that the observer computes an estimate based on the incorrect decision and then flips it to the correct side after feedback. (**c**) Hypothesis 2: Memory representation is preserved after the decision and the conditioned prior is updated upon receiving feedback. Model 2a (Flip decision) posits that the observer combines the memory representation and the updated prior to perform a Bayesian estimate.

We performed two psychophysical experiments similar in design to previous experiments ^7,8,21,10^. In both experiments, subjects were briefly presented with a visual orientation stimulus; they then had to first make a judgment whether the stimulus’ orientation was clockwise (cw) or counterclockwise (ccw) of a reference orientation before providing an actual estimate of the orientation by adjusting a cursor. Different from previous experiments, however, we provided subjects with correct feedback on their categorical judgment before they performed the estimate. Note, that feedback was delivered with a delay of 500 ms after subjects’ category judgments, thus providing time for the judgments to potentially update stimulus information in working memory. Subjects were instructed to consider the feedback when providing their estimates, which they always did. Both experiments included stimuli with different levels of sensory uncertainty, and only differed in that in Experiment 1 the range of stimulus orientation was symmetric around the reference whereas it was asymmetric in Experiment 2.

## Predictions

Both hypotheses make identical predictions for trials where subjects’ categorical judgment was correct. Given previous results, we expect subjects’ behavior to be well described with the self-consistent observer model ^10,20^. However, predictions differ for trials where that judgment was incorrect and subjects had to change their categorical assessment. Hypothesis 1 assumes that information about stimulus orientation and the categorical judgment are jointly and inseparably represented in working memory (Fig. 1b). Specifically, we make the assumption that after the categorical decision, the memory representation is updated to reflect the trial posterior distribution. According to the self-consistent observer model the posterior is zero for all stimulus orientations that are not in agreement with the categorical judgment ^10,20^. Thus, when receiving feedback indicating that the categorical judgment was incorrect, the observer is left with having to generate an estimate of the stimulus orientation based on the resulting posterior and the feedback. Here, we consider two different ways the observer may perform estimation with the modified memory representation. Model 1a assumes that the observer only relies on the prior distribution over orientation that corresponds to the correct stimulus category (feedback) because the posterior does not provide any information for orientations on the correct side of the reference. As a result, the model predicts an orientation estimate that represents the mean of the prior corresponding to the optimal estimate under mean squared-error loss. The estimate is on average the same across all stimulus orientations and levels of sensory uncertainty, and its variance is only depending on potential motor noise. Model 1b is a heuristic model that assumes that in case of an incorrect categorical judgment the observer performs an estimate based on the stored posterior but then simply flips its value around the reference. This model predicts estimate mean and variances that depend on stimulus orientation and uncertainty, but will not accurately reflect any potential asymmetry in the category priors around the reference.

Under Hypothesis 2, sensory information and the categorical judgment are independently represented in working memory (Fig. 1c). We consider a model (Model 2a) where the observer can flexibly re-combine the sensory information stored in working memory (i.e. likelihood) with the correct category prior in order to estimate stimulus orientation. This model is equivalent to the self-consistent observer model ^10^ under the assumption that the observer has the flexibility to simply flip the categorical judgment. Note, nothing in the inference process is affected by the observer’s initial, incorrect categorical judgment. Later in the paper we will also test a slightly modified version of this model (Model 2b).

We evaluated the two general hypotheses based on a particularly strong form of cross-validation: We fit only the data representing correct trials (i.e. subjects provided a correct categorical judgment) and then used the fit parameter values to predict estimation behavior in the incorrect trials for all above models. Models are fully constrained by the parameters of the self-consistent observer model, and thus the model comparison is parameter-free. See Methods for more details.

### Experiment 1: Results and model analysis

Five subjects (S1-5; S1 non-naïve) participated in Experiment 1. In each trial, an array of small, oriented line segments was briefly (500 ms) presented (Fig. 2a). After the stimulus disappeared, subjects had to indicate whether stimulus orientation was cw/ccw relative to a reference orientation. Auditory feedback indicated whether the categorical judgment was correct or not. Subjects were instructed that feedback was always valid, which it was. After receiving feedback, subjects had to report perceived stimulus orientation by adjusting a joystick.

**Figure 2:**
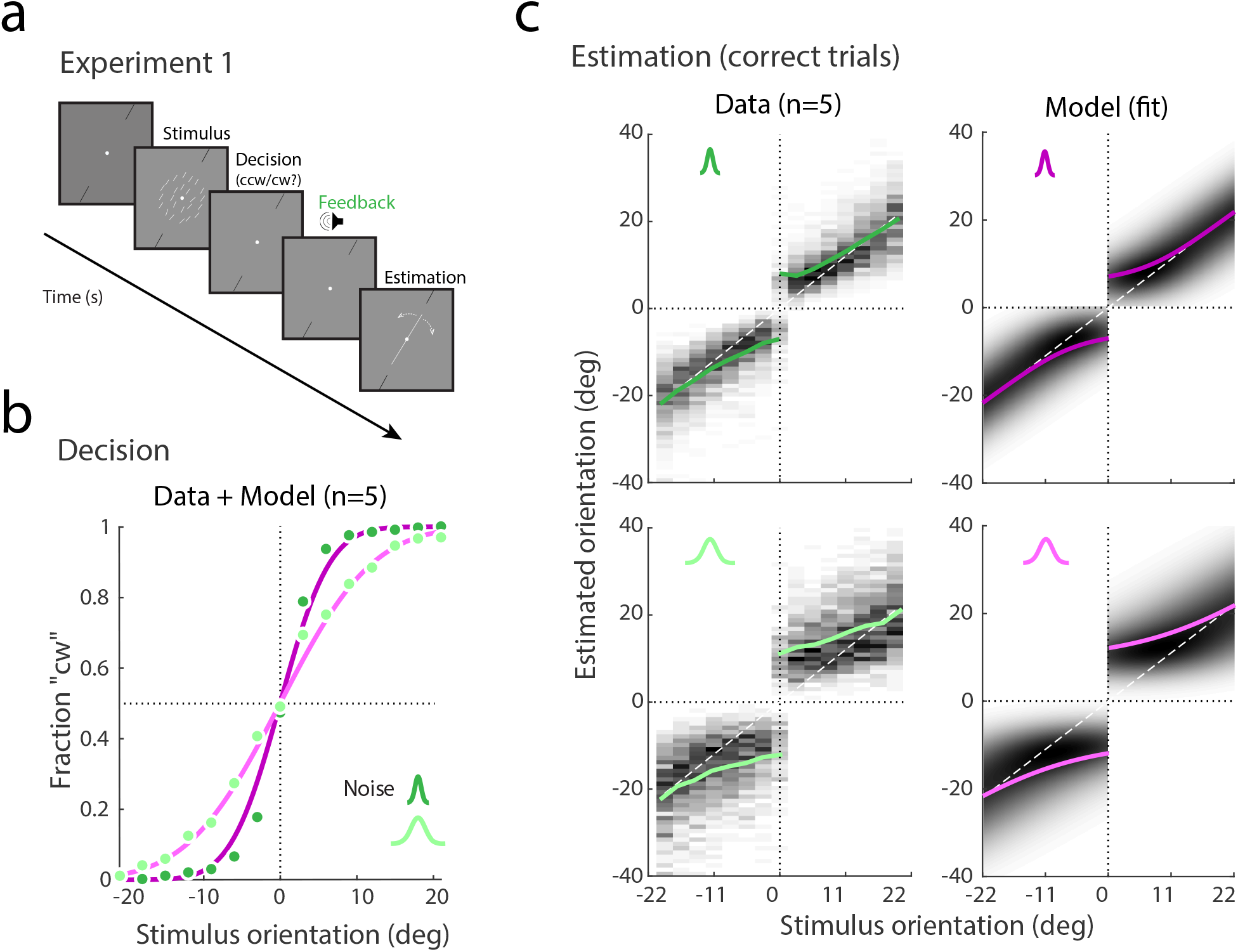
Experiment 1: data and model fits for correct trials. (**a**) The stimulus consisted of an array of line segments whose orientations were sampled from a Gaussian centered at a stimulus orientation. Stimuli had one of two levels of noise corresponding to different widths of the Gaussian (*σ_s_* := {3, 18} deg). Stimulus orientation was uniformly distributed within a range *±* 21 deg around the reference orientation. Subjects first reported whether the stimulus orientation was cw/ccw of the reference orientation (indicated by two black lines). 500 ms after reporting their decision, they received auditory feedback indicating whether it was correct or incorrect. Instructed to take feedback into account, subjects then provided an estimate of stimulus orientation by adjusting a probe line. (**b**) Subjects’ categorical judgment data (green circles; combined subject, n=5) and the self-consistent observer model fit (purple lines). Darker shades represent data/fit for lower stimulus uncertainty. As expected, the slope of the psychometric function is steeper for lower noise level. (**c**) Estimate distributions of the combined subject and the predicted distributions of the model fit (colored lines represent mean estimates). Subjects show the characteristic repulsive bias away from the reference, confirming the results of previous studies ^7,8,10^. Note that the model was jointly fit to the decision and estimation data. See Supplementary Fig. S1 for individual subjects’ data.

Figures 2b and c show the categorical judgment and estimation data for the combined subject (pooled trial data across all subjects). Estimate distributions (Fig. 2c) only reflect correct trials. Data confirm previous results showing the same characteristic bimodal distributions: estimates are biased away from the reference orientation with biases being larger for larger stimulus uncertainty and stimulus values closer to the reference ^7,8,10^. The figures also show the predictions of the best fit self-consistent observer model. Consistent with our previous findings the model well accounts for both the psychometric function of the categorical judgment as well as the accurate bimodal distributions of orientation estimates ^10^.

Next, we predicted the estimate distributions for incorrect trials for all models based on the fit model parameters (shown in Fig. 4a). Figure 3 shows the predictions together with the experimental data (combined subject). The data reveal a few noteworthy characteristics. First, subjects’ orientation estimates were on average relatively constant and independent of the true stimulus orientation although the estimates and their variances were substantially larger in the high stimulus noise condition (Fig. 3a). Second, the distributions for any given stimulus orientation are long-tailed away from the reference orientation. While all models predict relative constant average estimates, Model 1a (prior only) predicts estimate distributions that are identical for both stimulus noise levels and are not long-tailed (Fig. 3b). However, Model 1b (flip estimate) and 2a (flip decision) capture these key characteristics of the data well, although Model 2a tends to underestimate the magnitude of the estimates. Finally, while approximately constant, there is a slight trend in subjects’ estimates to increase in magnitude for stimulus orientations increasingly away from the reference. Model 1b predictions show the opposite trend.

**Figure 3:**
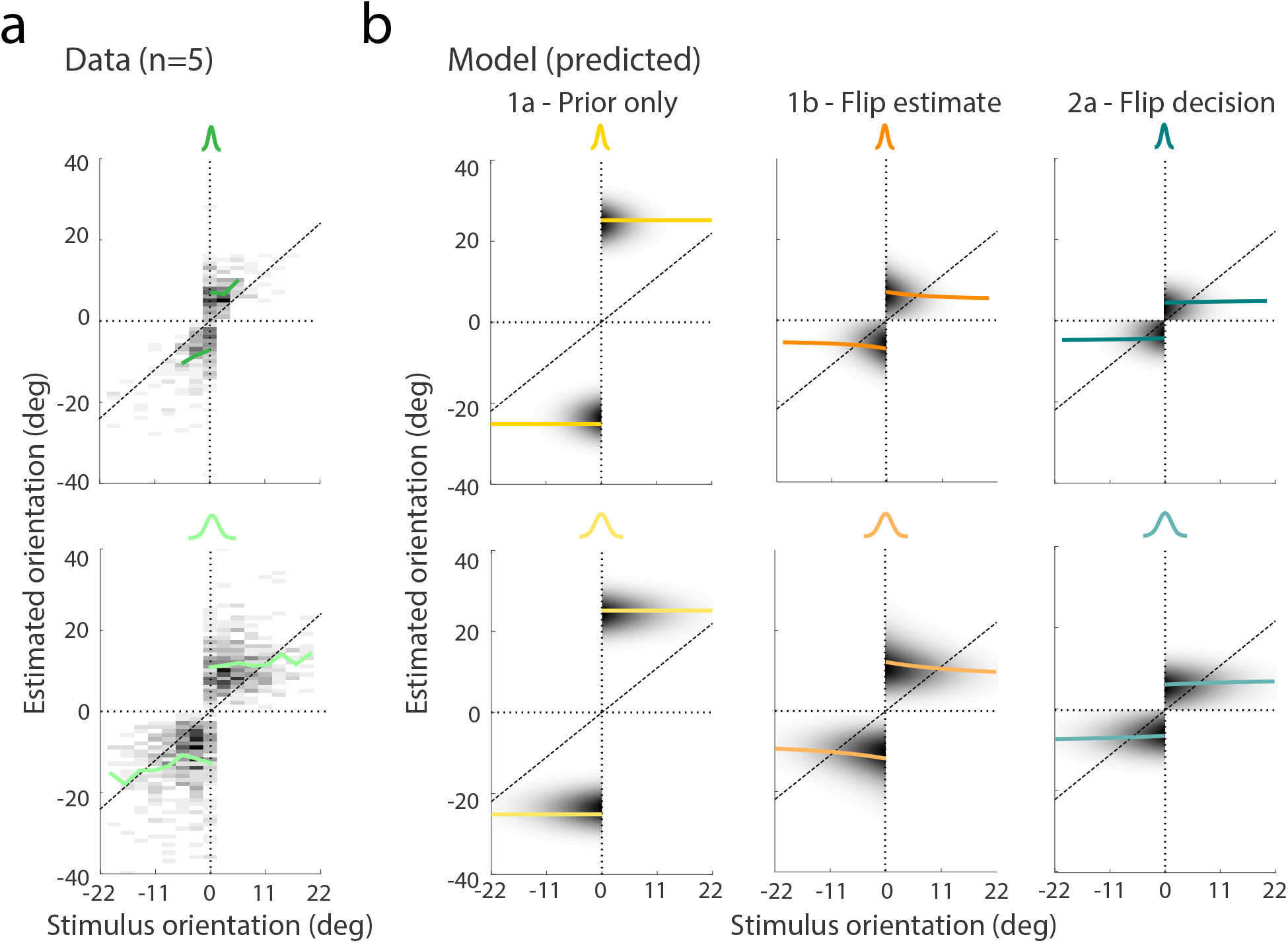
Experiment 1: data and model predictions for incorrect trials. (**a**) Distribution of subjects’ estimates in incorrect trials. Each row corresponds to a stimulus noise level (top: low noise). For stimulus orientations further away from the reference, the data get sparser because subjects made less incorrect decisions. Mean estimates (green lines) are approximately constant on each side of the decision boundary with magnitude and variance larger for higher stimulus noise. Also, distributions are long-tailed away from the reference orientation (see also Supplement Fig. S1) (**b**) All models correctly predict estimate distributions with approximately the same mean and variance independent on stimulus orientation. For Model 1a (prior only), however, the predicted mean corresponds to the mean of the prior range and the variance is only determined by the motor noise. This is not in agreement with the data. In contrast, the remaining two models (1b and 2a) correctly predict larger estimation means and variances for higher stimulus noise as well as long-tailed distributions. Both models provide a good account of the data although Model 2a tends to underestimate subjects’ actual estimates especially in the high noise condition, while Model 1b predicts slightly decreasing estimate magnitudes with increasing stimulus orientations away from the reference, which is somewhat against the overall trend in the data. See Supplementary Fig. S2 for individual subjects’ data.

**Figure 4:**
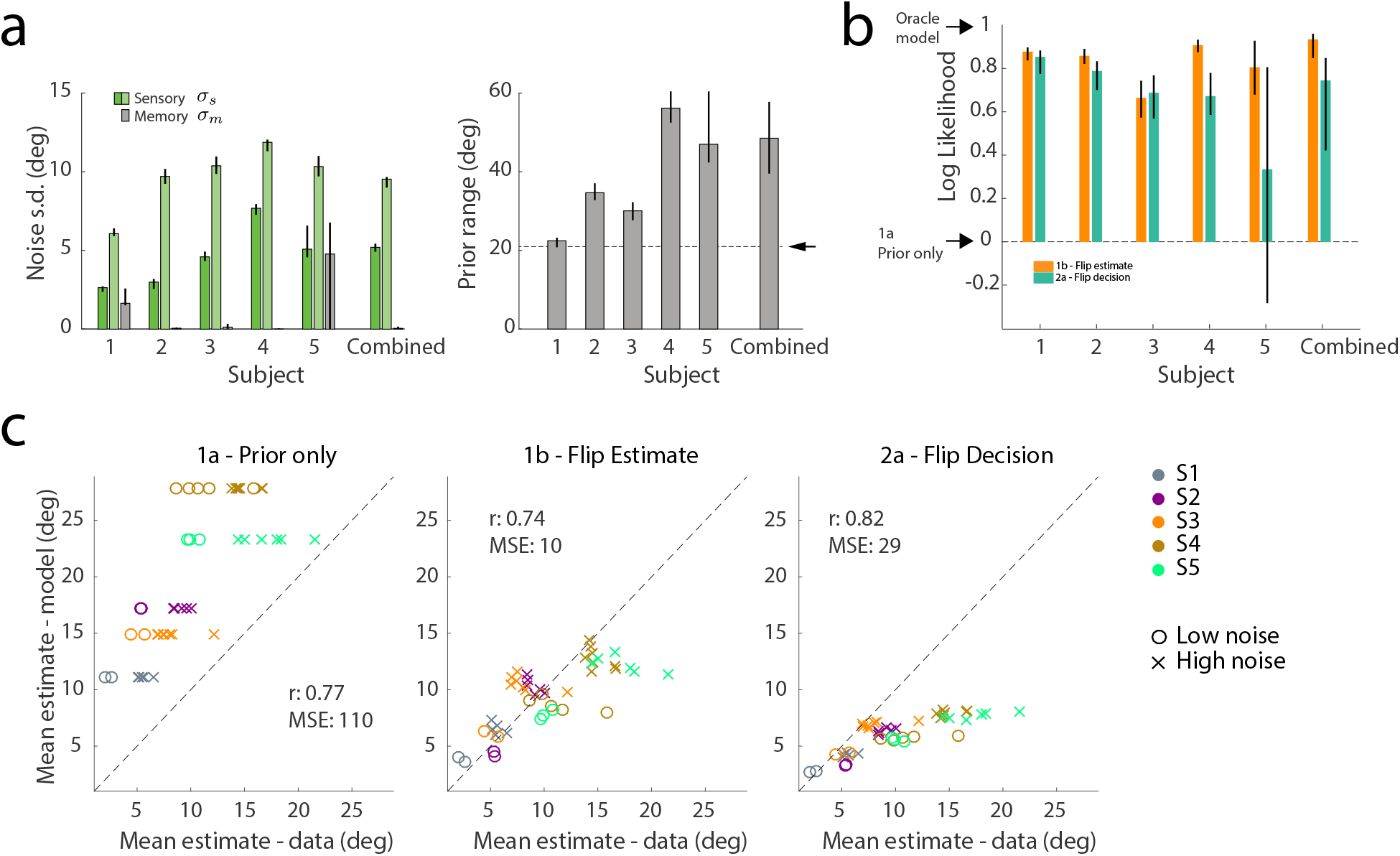
Experiment 1: individual model parameters and prediction performances. (**a**) Fit noise levels and prior range of individual subjects. While there is substantial inter-subject variability, the range of parameters values is consistent with our previous study ^10^. Also all subjects tend to overestimate the true stimulus range (*±* 21 deg from the reference orientation, arrow). (**b**) Log-likelihood comparison between the models. Values are normalized relative to the log-likelihood values of Model 1 (prior only) and the Oracle model (i.e. the likelihood value based on the empirical probabilities; see Methods for details). Model 1a performs substantially worse than the other models for all subjects. However, log-likelihoods for Models 1b and 2a are not statistically different except for subject 4. All errorbars in (a) and (b) indicate 95% confidence intervals computed over 200 bootstrapped samples. (**c**) Scatter plots comparing the mean estimates between models and data for every stimulus orientation (absolute values). Each point corresponds to one experimental condition (stimulus noise and orientation) of one subject. Although all models can roughly explain the trend in data, the prediction of Model 1a is substantially off from the data. Compared to Model 1b, Model 2a has a higher correlation but is worse when comparing mean-squared error (MSE) between model and data mean. Model 2a tends to underestimate the magnitude of subjects’ estimates on the higher end.

We complemented the above qualitative model comparison with a quantitative analysis of individual subjects’ data. As with the combined subject, we first fit individual subjects’ data for correct trials and then used the fit model parameters to predict estimate distributions for the incorrect trials. Figure 4a shows the fit model parameters for all individual subjects (and the combined subject). The fit parameter values are consistent with those in our previous study ^10^. In particular, subjects showed the same tendency to over-estimate the overall stimulus range to be approximately twice its size. All models can predict the full distribution of estimates (i.e. they are observer models). Thus, we computed the likelihood of each model’s prediction given the experimental data. Figure 4b shows the likelihoods for each subject normalized to the range set by the likelihood for Model 1a and the “Oracle model”, i.e. the likelihood based on the empirical probabilities of the data. The comparison is consistent across all subjects and reflects our initial, qualitative comparison. Model 1a (Prior only) is significantly worse than the other models. Although Model 1b (Flip estimate) performs best for most subjects, it is not statistically different from Model 2a (Flip decision) except for subject S4.

In order to quantify which model better captures the characteristic overall bias patterns in the data, we performed a second analysis. We compared the mean estimates for each stimulus condition (stimulus noise and stimulus orientation) for each subject with the corresponding predictions of the different models. Figure 4c shows this comparison together with the calculated correlation and mean squared-error (MSE) between model predictions and data. Although correlations are high for all models (0.74-0.82), the MSE is substantially higher for Model 1a than the other models. Model 2a (Flip decision) performs better than Model 1b (Flip estimate) in terms of correlation but not MSE. That accords with our above qualitative comparison in which Model 2a better predicts the trend in data but underestimates the magnitude in high-noise condition.

The experimental and analytical results of Experiment 1 suggest that Model 1a is not an adequate description of subjects behavior. In particular, subject’s working memory representation of the stimulus seems to retain at least some of the sensory information that was not in agreement with the initial categorical judgment. Also, this information must be stored in a way that is independent of a particular (high-level) categorical interpretation of the stimulus and thus can be flexibly recombined with the feedback information. However, the data are equally well explained by Model 1b and 2a. Thus we can not rule out that subjects directly store a biased estimate of the stimulus orientation in working memory and then just perform a mental mirroring at the reference orientation if feedback requires a change. Which is why we conducted Experiment 2.

### Experiment 2: Results and model analysis

Experiment 2 was similar to Experiment 1 except that the range of stimulus orientations was shifted to the cw side ([−12, 30] deg). Note that the stimulus distribution was still uniform on either side and the fraction of stimulus orientations being cw or ccw was still 0.5. If subjects were able to exploit this asymmetric stimulus distribution in form of prior knowledge then their estimates should reflect this asymmetry. However, Model 1b or 2a make opposite predictions for the sign of the asymmetry in incorrect trials, providing us with a qualitative criterion to disambiguate the two models.

Figure 5a outlines the structure of a single trial in Experiment 2. Subjects were extensively trained prior to the actual experiment in order to learn the asymmetric stimulus distribution. In addition, subjects were reminded of the actual stimulus range with a visual cue at the beginning of each trial. The rest of the trial design was identical to Experiment 1. Seven subjects participated in the experiment of which only one (non-naïve subject S1) also participated in Experiment 1.

**Figure 5:**
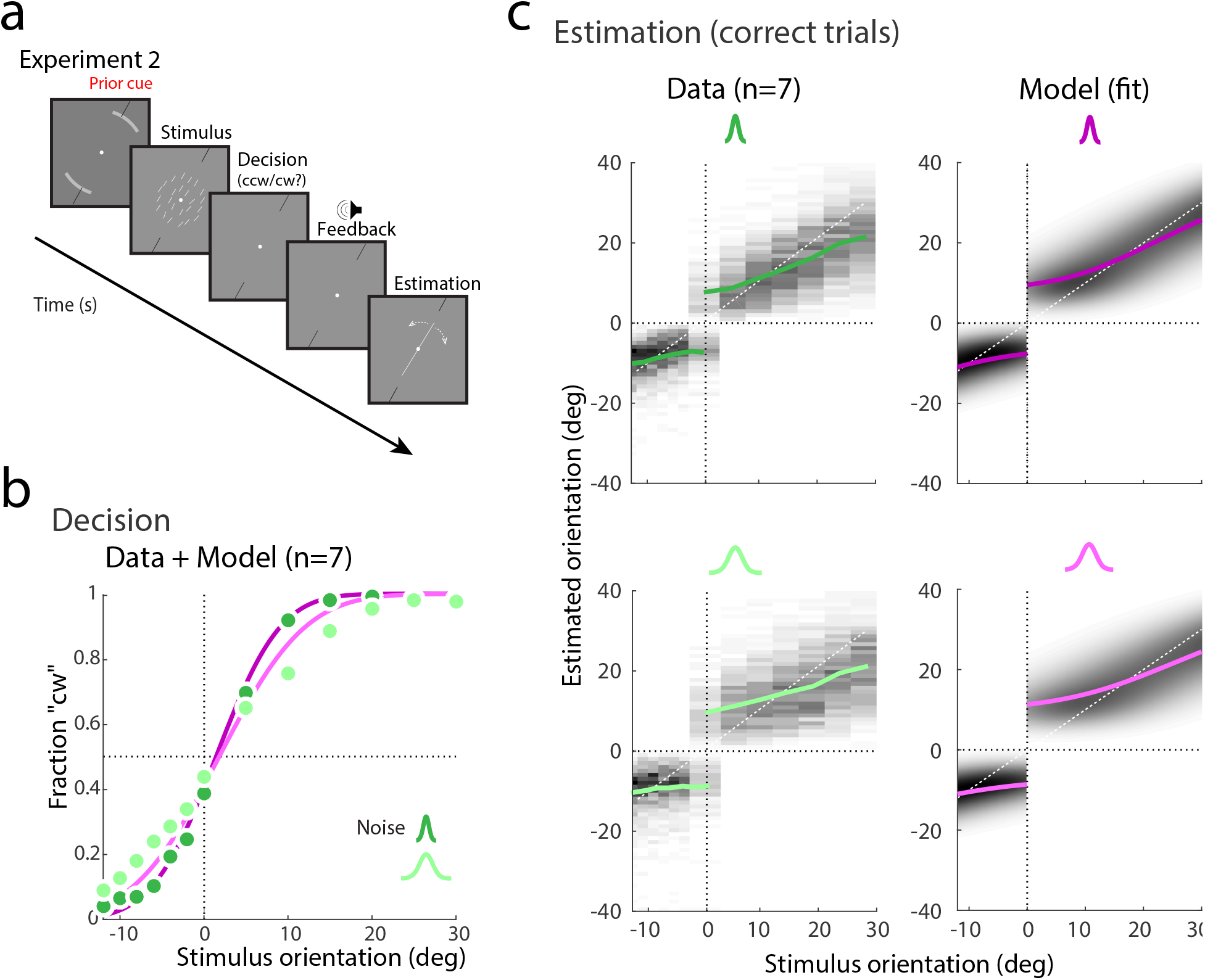
Experiment 2: data and model fits for correct trials. (**a**) Experiment 2 was similar to Experiment 1 except that the stimulus range was asymmetric around the reference ([−12, 30] deg, positive values indicating cw orientations relative to the reference). Subjects were reminded about the asymmetry at the beginning of each trial by explicitly showing the actual stimulus range (gray arcs). Stimulus noise levels corresponded to two different widths of the stimulus array distribution (*σ_s_* := {6, 18} deg). (**b**) Psychometric curves representing subjects’ categorical choice behavior (combined subject, n=7). Green circles indicate the data and purple lines indicate the fit with the self-consistent observer model. Darker shades represent conditions with lower stimulus uncertainty. Note that subjects’ decision probability is not exactly 0.5 for stimuli at the reference orientation. We allowed the category prior probability to be a free parameter in the model although fit values were close to 0.5 (see Fig. 6a). (**c**) Measured estimate distributions and the predictions of the fit self-consistent model for correct trials only (combined subject). Estimates on the ccw side are closer to the reference and less repulsed. This is well captured by the model and thus can be explained as the effect of the asymmetric prior in the inference process. See Supplementary Fig. S3 for model fits to data of individual subjects.

Figure 5b shows data for the categorical judgment and the fit with the self-consistent observer model (all trials; combined subject). The closer the stimulus orientation is to the reference and the larger the stimulus uncertainty, the more incorrect judgments were made. Figure 5c shows the estimate distributions for each noise level together with the predictions of the self-consistent observer model (fit to the correct trial data only). Estimates have a larger repulsive bias away from the reference orientation for stimulus orientations on the cw side of the reference. This is particularly visible in the high noise condition and clearly indicates that subjects took the asymmetric stimulus range into consideration when estimating stimulus orientation. The model well captures this characteristic suggesting that the asymmetry in estimation bias is fully explained by a Bayesian inference process with asymmetric prior. Fit prior parameters confirm this, showing values that are close to the true orientation range (Fig. 7a).

**Figure 6:**
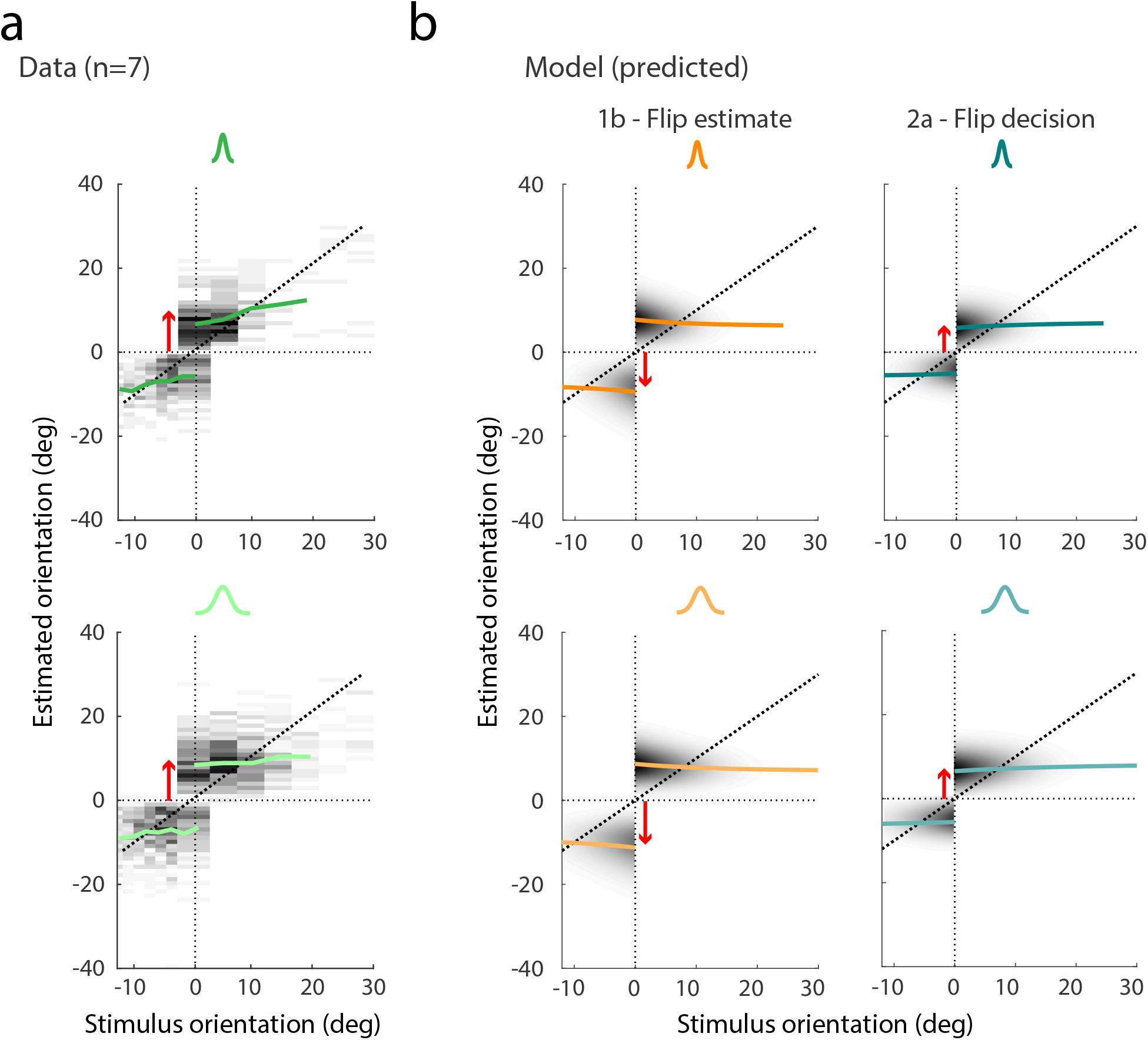
Experiment 2: data and model predictions for incorrect trials. (**a**) Distributions of subjects’ estimates in incorrect trials (combined subject). As in Experiment 1, estimates are approximately constant and independent on stimulus orientation. However, the distributions show a clear asymmetry with estimates on the cw side being further repulsed from the reference orientation than on the ccw side. This pattern suggests that subjects were able to flexibly change and incorporate the correct prior in their estimate according to feedback. (**b**) While Model 2a predicts the same asymmetry as seen in the data Model 1b predicts the opposite pattern. Red arrows indicates the side with larger repulsive biases. See Supplementary Fig. S4 for data of individual subjects.

**Figure 7:**
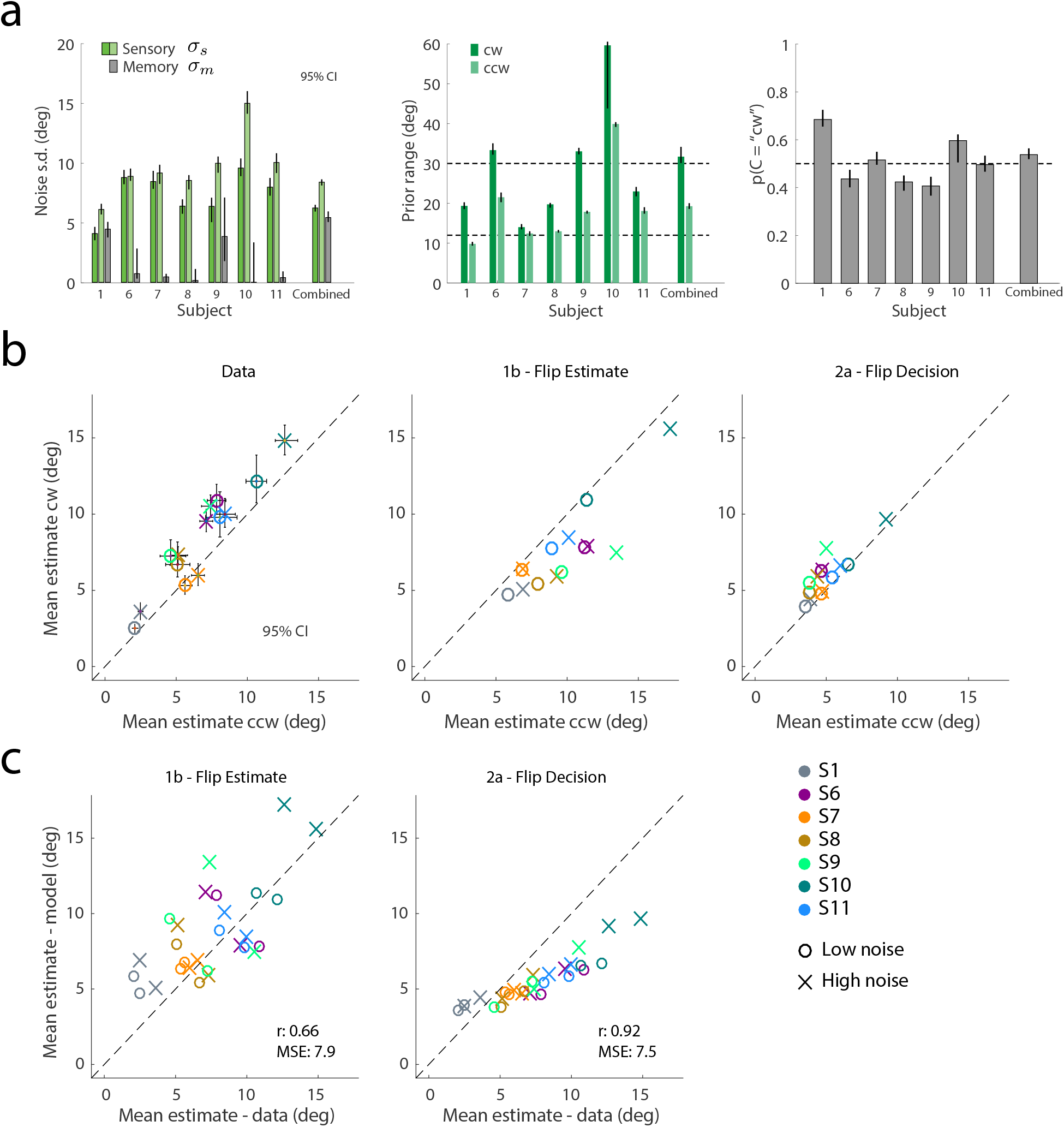
Experiment 2: individual model parameters and prediction performances. (**a**) Fit sensory (*σ_s_*) and memory (*σ_m_*) noise parameters of individual subjects. The fit prior range is consistently higher on the cw side for all subjects. We allowed the categorical prior *p*(*C* = “cw”) to be a free parameter (fixed in Experiment 1). On average the values are close to the statistically correct value of 0.5. (**b**) Mean estimates (averaged across stimulus orientations) in incorrect trials are larger on the cw side for all except one subject (S7). That pattern is consistent with the predictions of Model 2a but not of Model 1b: Flipping the estimate leads to larger estimates on the ccw side. Note, estimates are consistently larger for larger stimulus uncertainty. (**c**) Model 2a performs better than Model 1b in terms of both correlation and MSE. However, Model 2a predictions show the same tendency to underestimate estimation magnitudes as in Experiment 1.

For incorrect trials, observing the same asymmetry as in correct trials with larger estimates on the cw side would suggest that subjects can *flexibly switch* their categorical prior (to the one corresponding with feedback) and re-combine it with stored sensory information when estimating stimulus orientation. This is indeed what we found (Fig. 6a): As in Experiment 1, subjects’ estimates in incorrect trials are approximately independent of stimulus orientation but show the same asymmetry as in the correct trials; the average estimate is larger on the cw than on the ccw side of the reference (red arrow). Models 1b and 2a make markedly different predictions (Fig. 6b). While Model 2a (flip decision) follows the data, Model 1b (flip estimate) predicts a flip in asymmetry such that estimates on the ccw side are further away from the reference orientation. This clearly speaks against Model 1b.

We also performed a quantitative model analysis for data of individual subjects. The fit model parameters (correct trials only) are presented in Fig. 7a. Noise parameters are consistently larger for the high-noise stimulus. Fit prior parameters suggest that the behavior of all subjects, to various degrees, reflects knowledge of the asymmetric stimulus range and, on average, the correct category prior of 0.5. Figure 7b shows subjects’ mean estimates in incorrect trials and the corresponding predictions of Model 1b and 2a. Like for the combined subject, most subjects (6 out of 7) have mean estimates that are significantly larger on the cw than on the ccw side. Also, subjects’ estimates are consistently larger in the high noise condition. The predictions of Model 2a well captures this pattern while Model 1b predicts a mirrored pattern where estimates are larger on the the ccw side. The predictive log-likelihood, MSE, and correlation comparison between the two models further confirms that Model 2a is a better predictor for the estimation pattern in incorrect trials than Model 1b (Fig. 7c and Fig. 9b).

**Figure 8:**
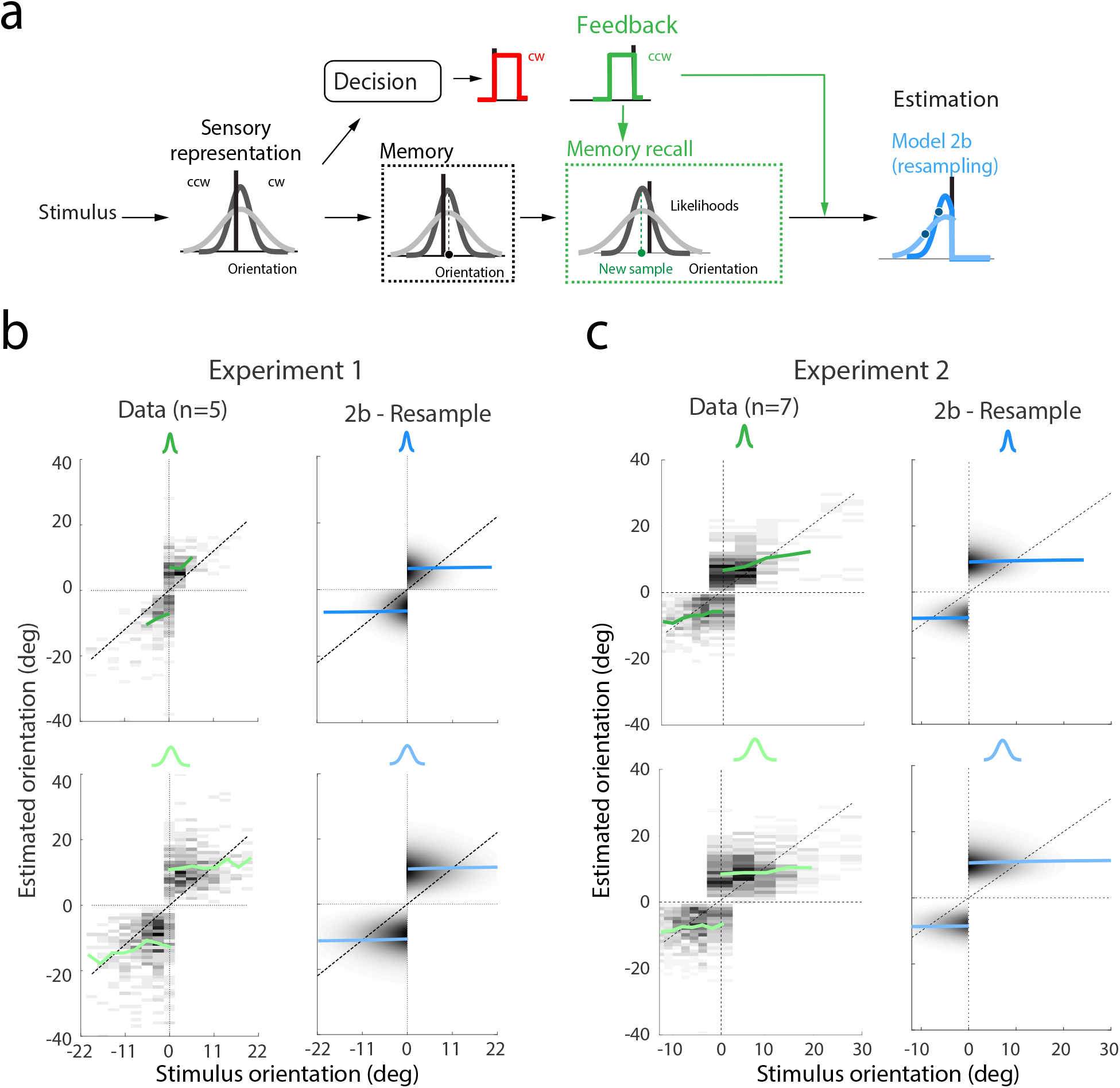
Conditioned memory re-sampling: predictions. (**a**) Model 2b is identical to Model 2a (flip decision) except that it assumes that the observer re-samples sensory measurements from working memory until a sample is obtained that lies on the correct side of the reference orientation (i.e. is consistent with the feedback in incorrect trials). A new likelihood function based on this sample is combined with the updated categorical prior to compute the posterior and, ultimately, the estimate. (**b**) Model’s prediction for Experiment 1 (combined subject). The model captures the overall shape, magnitude, and skewness of subjects’ estimate distributions. (**c**) Model’s prediction for Experiment 2. The model correctly predicts two important features of the data: the overall shift towards the cw side due to the asymmetric distribution, and the average magnitude of the estimates on each side. See Supplementary Figs. S6–S10 for more details.

**Figure 9:**
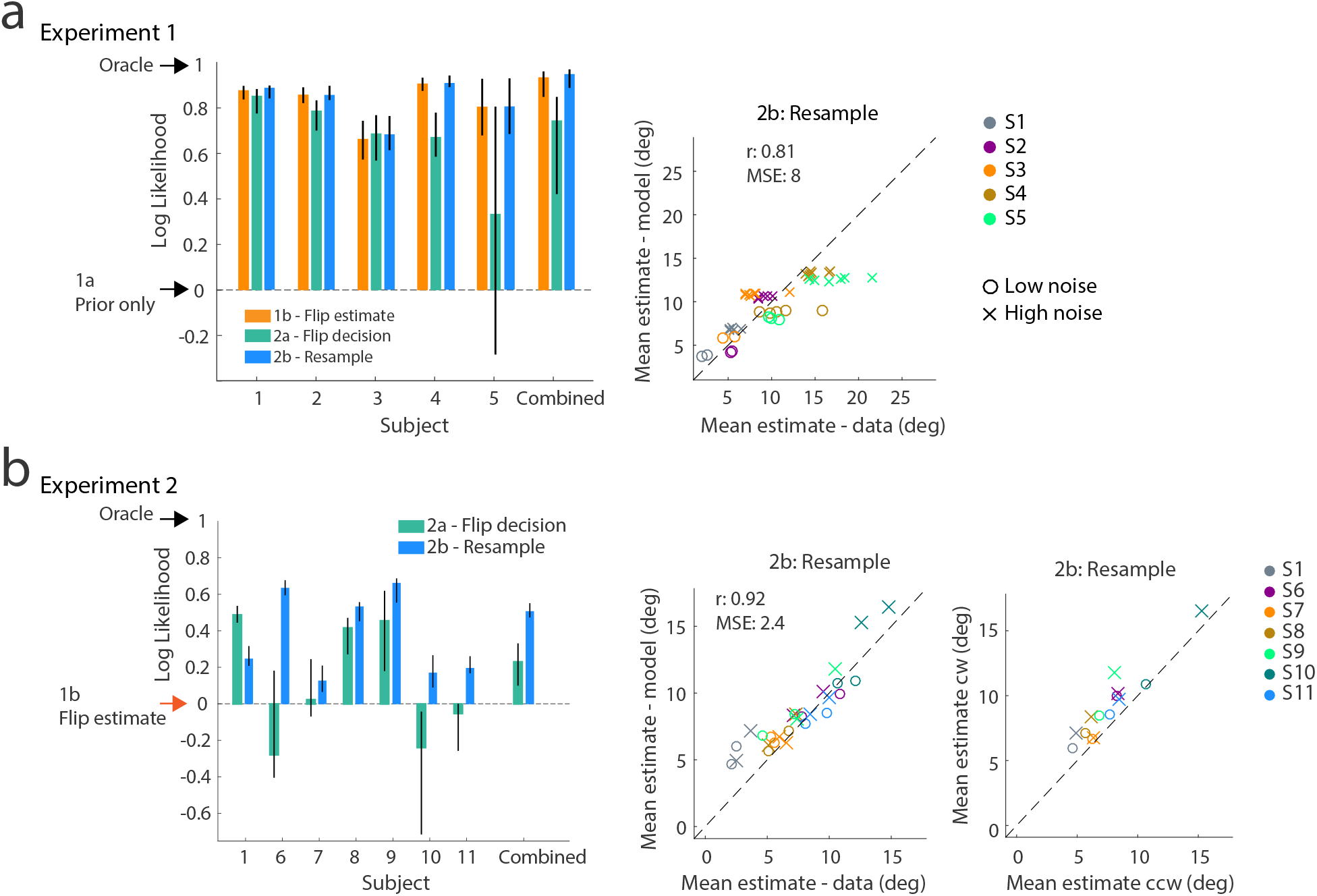
Conditioned memory re-sampling: model comparison. (**a**) Normalized log-likelihood comparison for incorrect trial data in Experiment 1. Model 1b, 2a, and 2b perform similarly. However, in terms of correlation and MSE, Model 2b outperforms the other models (see Fig.4c for comparison). (**b**) Comparison for incorrect trials in Experiment 2. Although close, the likelihood of Model 2b is consistently higher than the likelihood of Model 2a except for S1 (non-naïve). The correlation and MSE metrics also indicate that predictions of Model 2b are better than of Model 2a (see Fig.7c for comparison). The re-sampling model also correctly predicts mean estimates higher on the cw side compared to the ccw side (see also Fig. 7b). Note that all model predictions are parameter free and are based on the parameter values of the fit self-consistent observer model to the data in correct trials. Errorbars indicate 95% confidence intervals computed over 200 bootstrapped samples

Based on the results of Experiment 2 we can safely rule out that subjects estimate in incorrect trial reflect mirror copies of estimates made under their initial categorical interpretation. Their initial (incorrect) categorical judgment did not reduce sensory information to a point estimate representation in working memory. Rather, subjects seem to have maintained the initial sensory representation and its uncertainty in a way that allowed them to flexibly recombine it with the new, correct category assignment in an inference process that considers both stimulus uncertainty and priors. This is an important finding that has implications for understanding working memory function beyond the current experimental task. Nevertheless, across both experiments predictions of Model 2a slightly but consistently deviate from the data: the magnitudes of the predicted estimates are generally smaller than the data, which is particularly noticeable in high stimulus noise conditions (see Fig. 4c and Fig. 7d). As a result, we next consider a modification to Model 2a that further improves the quality of the predictions.

### Consistent re-sampling of memory representation

Model 2a represents the self-consistent observer model ^10^ where inference is conditioned on the feedback rather than the observer’s initial incorrect categorical judgment (see Fig. 1b). Sensory information is assumed to be independently stored in working memory in form of a measurement sample *m* and the associated uncertainty (i.e. the likelihood function). When inferring stimulus orientation, sensory information is recalled from memory by drawing a sample *m_m_* from the memorized measurement and updating the likelihood function accordingly (see Methods).

For the new Model 2b we now assume that memory recall is not a random sampling process (e.g. Gaussian around *m* with *σ_m_*) but constrained to provide samples that are consistent with the top-level categorical interpretation. This represents a slightly modified self-consistent observer model where the categorical judgment not only acts as a top-down prior but also as a constraint on memory recall (see Fig. 8a). The modification does not lead to any noticeable difference in the estimate distributions for correct trials (see Supplementary Figs. S6–S8). For incorrect trials, however, re-sampling predicts estimate distributions that are more repulsed from the reference orientation. Figure 8b and c show the predicted distributions together with the data for both Experiment 1 and 2, respectively. To maintain consistency, we used the re-sampling scheme both in the fit to correct trials and the prediction of incorrect trials. Figure 9 further quantifies the improvement in predictions. While making similar predictions, Model 2b is consistently better than Model 2a in accounting for individual subjects’ data (with the exception of non-naïve S1) for both experiments and across all performance metrics (log-likelihood, correlation, and MSE).

Re-sampling takes time in particular if the measurement *m* is far on the incorrect side of the reference and thus, on average, many samples have to be drawn before one is acceptable. An immediate prediction of the re-sampling model is that the estimation process takes longer in incorrect than in correct trials. We analyzed the time it took subjects to perform the estimation task in Experiment 2 (Fig. 10a). We defined subjects’ response time (RT) as the time between subjects’ confirmation of their decision and confirmation of their estimate. Consistent with the model’s general prediction, subjects had substantially larger RTs for incorrect than correct trials (difference: 430 ms, correct: 2530 ms, incorrect: 2960 ms). In addition, we found a consistent trend that RTs are larger for high than for low stimulus noise conditions. We measured increases in RTs of Model 2b in relative time-units that were proportional to the numbers of re-samples required before a sample is acceptable (simulation). Model RTs show the same relative increase for incorrect trials, as well as the trend of predicting slightly larger RTs for high noise stimulus conditions (Fig. 10b). Note that the latter is true for both correct and incorrect trials.

**Figure 10:**
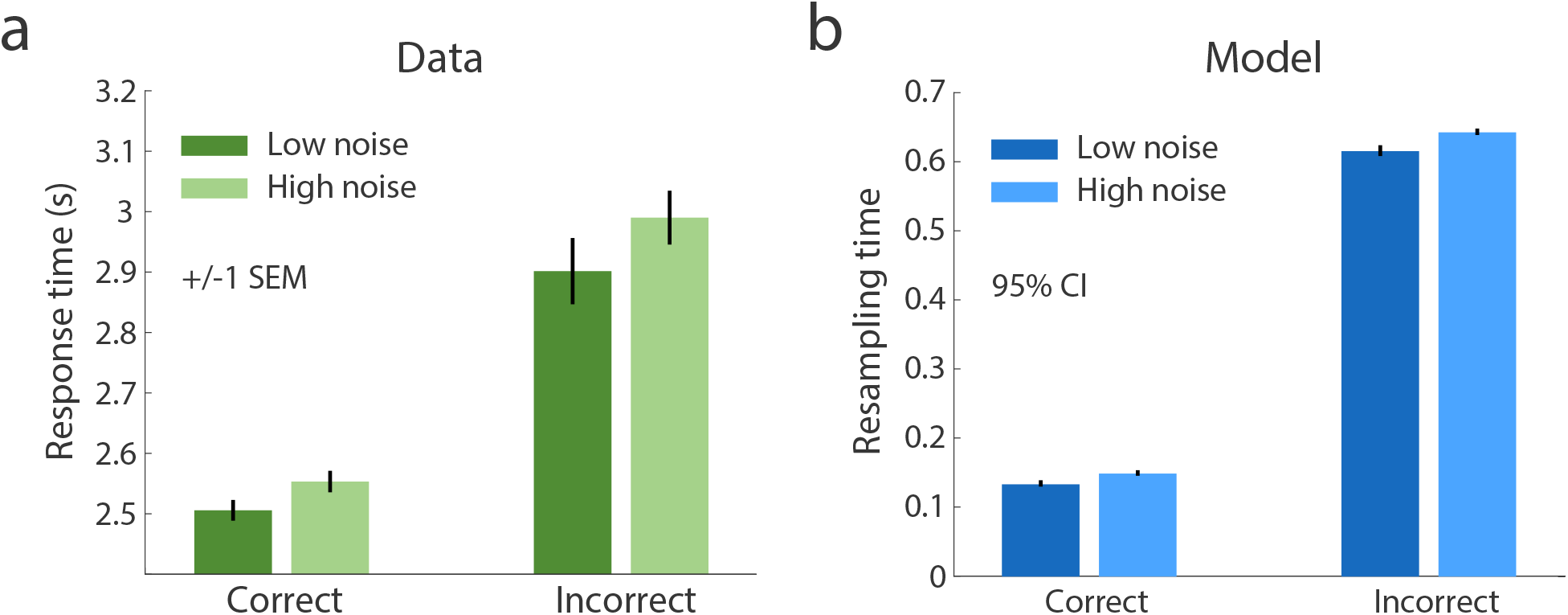
Response time (RT) for subjects and the conditioned re-sampling model. (**a**) Subjects’ RTs in the estimation task (Experiment 2). RTs are measured as the time between confirming the decision and confirming the estimate with a button press on the game pad. RTs are substantially higher in incorrect trials compared to correct trials (combined subject data). Also, RTs tend to be larger for higher noise condition. (**b**) Simulated increases in RTs of Model 2b. Increases in RTs are calculated as the relative number of samples necessary before a sample was acceptable (i.e. fell on the correct side of the reference). The simulated increases in RTs are qualitatively consistent with the data.

## Discussion

Previous studies have demonstrated how a high-level, categorical interpretation of a stimulus biases a subject’s subsequent recall of a low-level sensory feature of the stimulus. In this paper, we tested whether these biases directly reflect changes to the working memory representation of the stimulus or, alternatively, are the result of a perceptual inference process downstream from working memory. We ran two human psychophysical experiments that allowed us to measure the impact of subjects’ categorical assessments about the average orientation of a visual stimulus on their subsequent recall of stimulus’ orientation under different noise conditions and stimulus distributions. By providing feedback about their categorical judgments we were able to probe subjects’ memory recall in trials in which they were incorrect in their initial judgment and had to switch their high-level interpretation of the stimulus accordingly. Using a strong cross-validated model comparison, we found no evidence that subjects’ initial (incorrect) judgments lastingly modified working memory representation of the stimulus. Rather, we found that subjects were able to flexibly re-combine the original sensory information in working memory with the updated categorical interpretation in their perceptual recall process. Data from both experiments were remarkably well predicted by a modified self-consistent observer model that conditions recall and inference to be consistent with the provided feedback.

Our results have implications that generalize beyond the specifics of our experiments. Firstly, estimation biases were significantly different for high-compared to low-noise stimulus conditions. Given that noise conditions were randomly changed on each trial, this suggests that working memory representations of stimulus features include the representation of trial-by-trial sensory uncertainty. This confirms conclusions from previous studies ^24,25^, but furthermore also demonstrates that subjects can flexibly re-use this uncertainty information within a single trial. While our model reveals little about the physiological implementation of working memory, it is worth noting that it is formally identical to early Bayesian observer models in perception e.g. ^4^: Sensory information is represented by a trial specific measurement *m* and description of uncertainty centered on that measurement (i.e. a likelihood function). Inference is based on a recalled measurement that itself is a sample of *m* according to the memory noise distribution and a likelihood function that reflects this additional uncertainty together with the initial encoding uncertainty.

Secondly, the ability to flexibly re-combine sensory information with a different category interpretation are attributes shared with other models suggesting that humans can efficiently re-use parts of previous inference processes in subsequent inference steps ^26,27^. However, in contrast to our model these “amortized inference” models assume re-combinations at the level of entire posterior distributions, which cannot explain the bi-modal estimate distributions in the experimental data of the current and previous studies ^7,8,10^.

Also, the demonstrated flexibility of human subjects to re-combine information at different levels of the representational hierarchy has implications for our general understanding of visual working memory. While previous results suggest that working memory representations are tightly linked across different levels of a hierarchical stimulus representations (see review ^28^) our findings show that these links remain sufficiently flexible at the time of recall.

Thirdly, the model that best predicted subjects’ estimates in incorrect trials assumes that memory recall involves a feedback-consistent re-sampling process. Interestingly, such re-sampling is conceptually similar with ideas of retrospective shifts in attentional focus during memory recall ^29^: The stimulus used in our experiments consisted of an array of oriented line segments. Thus working memory representations likely reflect distributions rather than only the average orientation as we assumed for reasons of simplicity. Feedback can act as a retrospective attentional shift, boosting the accuracy of memory representations of those lines that are consistent with feedback, which then results in a bias in the recalled average representation towards the side consistent with feedback. Future work is needed to more thoroughly investigate the similarities between retrospective attention and the proposed re-sampling process within Bayesian self-consistent inference.

Re-sampling may not be limited to conditions where a subject is forced to change the categorical judgment but rather reflect an intrinsic mechanism of the type of hierarchical sequential inference tasks considered in our study. If so, it would require a modification of the self-consistent observer model ^20,10^. However, both the original and the updated model provide near-identical fits to data from correct trials in our experiments (see Supplemental Figs. S6–S10). It confirms that re-sampling has little effect on the estimate in trials in which subjects did not have to change their categorical interpretation of the stimulus. Thus we expect the updated model to provide an equally valid explanation of all the previous data sets ^7,8,10^ and therefore represent an improved implementation of the self-consistent observer model.

Finally, our results may provide new interpretations of some well-known bias phenomena in cognitive judgment and decision making. Hind-sight bias, for example, refers to subjects’ biases in recalling their initial assessment of an uncertain quantity upon receiving feedback ^30,31,32^. For example, subjects were asked to judge the probability of an event. After having received feedback on the correctness of their answer, subjects’ recall of their initial answer was significantly biased towards being more consistent with the feedback. This is generally consistent with our results. However, our results suggest that these biases are due to a conditioned inference process during memory recall rather than a direct, feedback-induced modification of memory representations ^33^. As such, hind-sight bias may be malleable or can even be eliminated by appropriate, measured feedback.

We end by proposing the following hypothesis: Visual working memory reflects an accurate reverberation of the original visual stimulus that, however, is flexibly interpreted at the time of recall. Interpretation is the result of a hierarchical inference process that allows to potentially incorporate additional or newer knowledge (e.g. feedback). As a result, the same memorized sensory information can be interpreted multiple times in different ways if necessary. Crucial, however, is that the interpretation is self-consistent such that a commitment at one level conditions inference at the rest of the hierarchy. Working memory recall is an active, reversible, but self-consistent inference process.

## Methods

### Experimental procedure

Eleven subjects (3 males, 8 females; subject S1 non-naïve) with normal or corrected-to-normal vision participated in the experiments. All subjects provided informed consent. The experiments were approved by the Institutional Review Board of the University of Pennsylvania under protocol #819634.

#### General procedure

Subjects sat in a darkened room. Stimuli were presented on a special purpose computer monitor (VIEWPixx3D, refresh rate of 120 Hz and resolution of 1920 x 1080 pixels) at a distance of 83.5 cm (Experiment 1) or 91 cm (Experiment 2) enforced via a chin rest. Screen background luminance was 40 cd/m^2^ and mean stimulus luminance was 49 cd/m^2^. Both experiments and data collection were programmed in Matlab (Mathworks, Inc.) using the MGL (http://justingardner.net/mgl) and Psychophysics Toolbox ^34^, and were run on an Apple Mac Pro computer with Quad-Core Intel Xeon 2.93 GHz and 32GB of RAM. Before each main experiment, subjects were extensively trained to get familiar with the task and to learn the stimulus distribution.

After training, each subject either completed 2100 trials in 3-5 sessions for Experiment 1 or completed 1820 trials in 3-4 sessions for Experiment 2. Each session lasted approximately 50 minutes. In total, each stimulus condition was repeated 70 times and consisted of 15 (Experiment 1) or 13 (Experiment 2) stimulus orientations and two noise levels. Subjects used a gamepad (Sony PS4 Dualshock) to indicate their categorical judgments by pressing a trigger button (left for ‘ccw’, right for ‘cw’). To report the perceived stimulus orientation, subjects adjusted a reference line (length: 5 deg) with the analog joystick of the gamepad and then pressed a button for confirmation. Subjects were instructed to always fixate whenever the fixation dot was present.

#### Experiment 1

Five subjects (S1-5) participated in Experiment 1. At the beginning of each trial, subjects were presented with a fixation dot (diameter: 0.3^*°*^) and a reference orientation indicated by two black lines (length: 3^*°*^, distance from fixation: 3.5^*°*^). We randomly selected the orientation of the reference from a uniform distribution ranging 0*°* to 180*°* (the full circle) in each trial. After 1 s, we presented an orientation stimulus consisting of an array of white line segments (length: 0.6^*°*^) that were arranged on two circles around at the fixation dot: the outer circle (diameter: 3.8^*°*^) has 16 line segments and the inner circle (diameter: 1.8^*°*^) has 8 line segments. Small random jitters (from *−*0.15*°* to 0.15^*°*^) were independently added to the x-y coordinates of each line segment. We sampled the orientation of each line segment from a Gaussian distribution whose mean (stimulus orientation) ranges from −21 (ccw) to 21 (cw) deg in steps of 3 deg with standard deviations (stimulus noise) 3 and 18 deg. Stimuli were presented for 500 ms. Subjects were instructed to indicate whether the average array orientation was cw or ccw of the reference orientation. Auditory feedback (100 % valid) was provided 500 ms after subjects’ response to indicate whether the categorical judgment was correct. Subjects then went on to report their perceived stimulus orientation. They were instructed that the feedback was always valid and should be taken into account when performing the estimation task. If subjects took longer than 4 s to provide the categorical judgment then the current trial was skipped and was moved to the end of the trial queue. At the end of each trial, a blank screen (mean luminance) was displayed with a duration randomly chosen from 300 ms to 600 ms.

#### Experiment 2

Seven subjects (S1, S6-11) participated in Experiment 2. The experimental design was identical to Experiment 1 except for the following differences: Stimulus range was [−12, 0] deg in steps of 2 deg for the ccw side and [0, 30] deg in steps of 5 deg for the cw side. We used different step sizes for the two sides to keep the total number of stimulus orientations on both sides the same. Also, subjects were reminded of the (asymmetric) stimulus range at the beginning of each trial for both training and main experiments (see Fig. 5a). Standard deviation of the low-noise condition was 6 deg compared to 3 deg in Experiment 1.

#### Training - simple motor task

Subjects were first trained to perform the estimation task. In each training trial, subjects first fixated their gaze on the fixation dot. For subjects participating in Experiment 2, we also displayed the decision boundary (like in the main experiment) and a gray arc indicating the asymmetric stimulus range during fixation. After fixation only the fixation dot remained and a white line (stimulus) appeared for 500 ms. Subjects then had to reproduce the white line by adjusting a line cursor with the analog joystick. After confirming their response by pressing a button, the original stimulus was displayed in green on top of subjects’ estimates. The decision boundary was uniformly sampled around the circle and the stimulus orientation was uniformly sampled in the same range as in Experiment 1 and 2, respectively. Subjects completed 450 trials (Experiment 1) or 325 trials (Experiment 2). We computed the standard deviation of subjects’ estimates and used that as the individual motor noise parameter in the model fits and predictions performed in our analysis.

#### Training - estimation task

After the first training session, a second training session was performed to accustom subjects with the stimulus estimation task. Trials were identical to the motor training except that we used the array stimulus in the main experiment instead of the single line stimulus. Subjects completed 100 trials (Experiment 1) or 150 trials (Experiment 2) for this training.

#### Training - decision task

The third training session was aimed at training the categorical judgment task. Stimuli were identical to estimation training but subjects were tasked to perform a categorical judgment (cw/ccw). Feedback was provided like in the main experiments. Subjects completed 900 trials (Experiment 1) or 200 trials (Experiment 2).

### Self-consistent Bayesian observer model

The key assumption of the model is that after making a categorical judgment, the observer fully conditions the subsequent estimation process on that subjective decision. The model is described in detail in Luu and Stocker ^10^, but for the reader’s sake we provide a summary description below.

### Decision task

A key assumption of the model is that the observer considers the stimulus in the context of a hierarchical generative model. Let *C* = {’cw’, ‘ccw’} be the category variable indicating whether the stimulus orientation is cw or ccw relative to a reference line and *θ* be the low-level stimulus feature representing the mean orientation of the line array. Furthermore, the model assumes that on each trial the observer forms a noisy sensory measurement *m* representing the stimulus orientation *θ*. Also, the observer knows the sensory uncertainty associated with that particular measurement *m* represented as a likelihood function *p*(*m|θ*). Assuming that the observer follows an optimal decision process, the likelihood function over *C* is given as

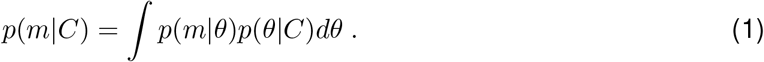

where *p*(*θ|C*) represents the observer’s prior expectation about the stimulus orientation on the two sides of the decision boundary. The observer then computes the posterior over the decision variable by combining the likelihood and the prior *p*(*C*):

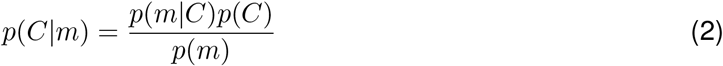

Given a symmetric loss function, the observer chooses the category with the higher posterior, thus

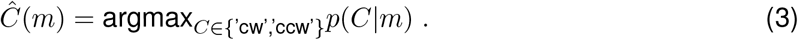

Note, that we assume the parameters of the likelihood function and the prior to be subjective, i.e., determined by the particular noise in the observer’s sensory process and their probably imperfect knowledge of the stimulus statistics. In that sense, “optimal” refers to the observer’s best knowledge and not to some sort of absolute ground-truth. We can obtain the model prediction of the psychometric function by marginalizing over the sensory measurement distribution:

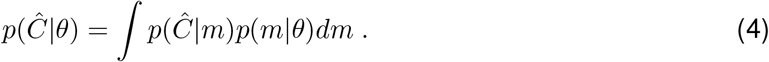

### Estimation task

After making the categorical decision, the observer’s task is to estimate the stimulus orientation. We assume that the sensory measurement *m* has been held in visual working memory since the stimulus disappeared. Further, we assume that at the time of estimation, *m* is recalled from memory with some additional uncertainty due to memory degradation. We refer to the recalled sample as *m_m_* and assume that memory degradation is described by the distribution *p*(*m_m_|m*). The distribution of memory samples for a given stimulus orientation is thus given as

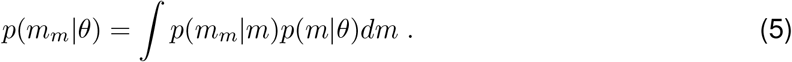

A fundamental aspect of the model is that the observer considers their own subjective categorical judgment as if it was factual. As a result, the preceding category judgment determines the prior distribution in the estimation task referred to as the conditioned prior 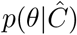. Thus the resulting estimate is determined by computing the posterior based on the likelihood function (Eq. (5)) and this conditioned prior,

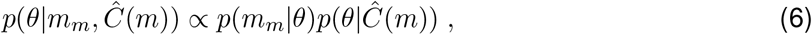

and then applying a particular loss function. Unless indicated otherwise, we assume an *L*_2_ norm loss, thus

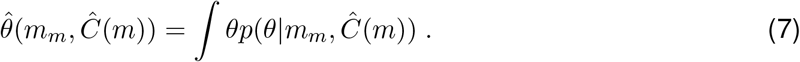

### Model predictions for incorrect trials

In testing their ability to correctly predict subjects’ estimates in those trials where their categorical judgment was incorrect, we can now formulate and compare different models that address the question whether categorical judgments lead to an update of working memory representations or not. Note that for correct trials, behavior is well described by the self-consistent observer model outlined above. For incorrect trials we considered the following models based on our two alternative hypotheses (Fig. 1) for how subjects’ initial categorical judgment potentially affects the stimulus representation in working memory.

### Hypothesis 1: categorical judgments update memory representations

We considered two potential ways the initial, incorrect categorical judgment 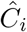 may have updated sensory working memory, either by updating the likelihood function to be consistent with the posterior (Model 1a) or directly replacing it with a point estimate (Model 1b), both according to the self-consistent observer model above conditioned on the initial choice 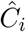.

#### Model 1a (prior only)

Updating the likelihood function based on 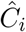 can be written as

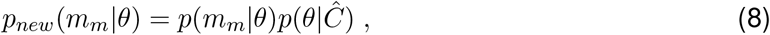

which erases sensory information that is inconsistent with the decision (i.e., the likelihood function is set to zero/uniform for that stimulus range). Thus, if feedback indicates that 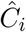 was incorrect, the posterior for those incorrect trials thus is equivalent to the prior distribution according to the feedback corrected judgment 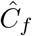, thus

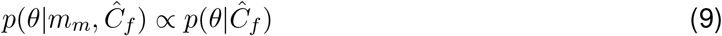

Assuming an *L*-2 loss function, the optimal estimate is represented by the mean of the posterior, i.e. the mean of the prior, thus

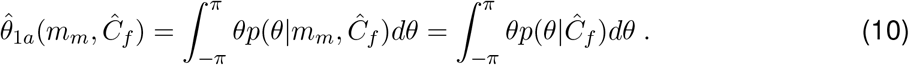

#### Model 1b (flip estimate)

After the initial (incorrect) categorical judgment, the observer immediately performs a self-consistent estimate 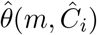 that then replaces the sensory representation in working memory. Then, according to feedback, the observer simply recalls this point estimate from memory (with noise) and flips it to lay on the correct side, thus reporting the estimate 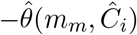.

### Hypothesis 2: sensory representation in working memory is preserved

Alternatively, we considered that sensory information is unaltered in working memory. Thus the information stored is equivalent to the original measurement *m* and some representation of uncertainty around that value (e.g. given by the likelihood function *p*(*m|θ*)). The general assumption is that the observer simply uses the category judgment 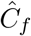 provided by feedback rather than the initial, incorrect judgment 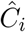 when performing the estimate. However, we consider two different versions of how working memory is recalled for this inference process.

#### Model 2a (flip d ecision)

M emory r ecall i s a s d escribed a bove l eading t o a l ikelihood function (Eq. (5)) that represents additional uncertainty due to memory degradation. Inference is according to the self-consistent observer model conditioned, however, on the feedback categorical judgment 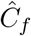, thus

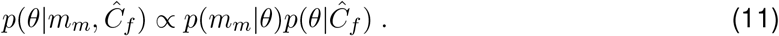

The estimate 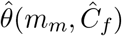 is then the mean of the posterior under *L*-2 loss function.

#### Model 2b (flip decision, re-sample)

This model is identical to Model 2a except that memory recall does not follow the distribution (5) but involves a conditioned sampling process 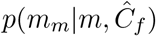 such that it only accepts memory recalls *m_m_* that are consistent with 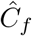. We assume that samples are repeatedly drawn until the sample lies on the correct side of the decision boundary.

For both models, the predictive estimate distribution are computed by marginalizing over the distribution of memory samples (or the re-sampled samples)

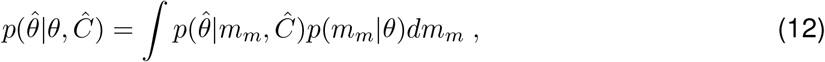

and then again marginalizing over the distribution of decision outcome (the psychometric function), thus

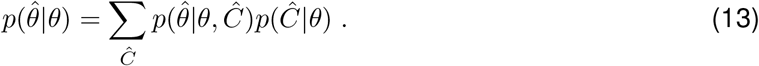

For all models, we convolve the corresponding predictive distributions with a motor noise kernel that was independently determined for each subject based on their individual motor training data (see above).

### Model specification

Note, model predictions were all parameter-free. Model parameters were based on fits of the self-consistent model on the correct trial data. The model contains the following parameters:

- The category prior *p*(*C*) is fixed to be 0.5 for Experiment 1 because of symmetry reasons. However, we leave this as a free parameter for Experiment 2 because the asymmetric stimulus range may induce a non-uniform subjective category prior (even though objectively *p*(*C* = ‘cw’) = *p*(*C* = ‘ccw’)).
- The prior distributions over stimulus orientation *p*(*θ|C*) are either symmetric (Experiment 1) or asymmetric (Experiment 2) around the decision boundary. More specifically, *p*(*θ|C*) is constant from the decision boundary up to a certain range on each side and then monotonically decreases to zero with a cosine roll-off. So the prior distribution is characterized by two parameters: the prior range *α* which is the distance from the decision boundary to half the magnitude of the uniform range and a smooth factor *β* which indicates how smooth the roll-off is. For the purpose of fitting and predicting the data, the role of smooth factor *β* is relatively small compared to the prior range *α*. Therefore, we only show the prior range in our plots of model parameters.
- The measurement distribution *p*(*m|θ*) is assumed to be Gaussian centered on the stimulus orientation *θ* and with standard deviation *σ_s_* proportional to the sensory uncertainty (variance in array). We consider *σ_s_* a free parameter for each subject and noise level that, however, only depends on the stimulus noise and is fixed across other experimental conditions.
- We assume working memory to be noisy such that the recalled sensory measurement *m_m_* represents a noisy sample of the original measurement *m*. Noise is modeled according to a Gaussian *p*(*m_m_|m*) centered on the original sensory measurement *m* with standard deviation *σ_m_*. The latter is assumed to be a free parameter for each subject but fixed for each subject across all experimental conditions.
- For Model 2b, the re-sampling distribution *p*(*m_m_|m*) is centered on the memory sample *m* and has the standard deviation 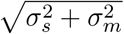.
- The motor noise is modeled as a Gaussian with standard deviation *σ*_0_ that was determined from the motor noise training data for each individual subject (see Supplementary Figs. S5).

### Model fit

We performed a joint fit to all trials of the decision task and correct trials of the estimation task by maximizing the likelihood of model parameters given the data:

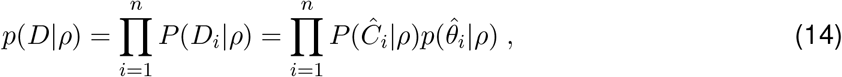

where *D* is the data, *ρ* is the parameters of the model, 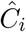 is subjects’ decision, 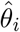 is subjects’ orientation estimate, *i* is the trial index and *n* is the number of trials. We used Nelder-Mead simplex optimization algorithm to minimize the negative log likelihood *−log*(*p*(*D|ρ*)). We ran the optimization routine thirty times starting at randomized initial parameter values to find the best parameter set.

## Data availability

The raw datasets will be provided as a text file upon the acceptance of the manuscript.

## Code availability

The code of computational models will be deposited in the public repository Github upon the acceptance of the manuscript.

## Acknowledgment

This work was supported by grants from the National Science Foundation (BCS-1350786 and IIS-1912232), and the University of Pennsylvania. Part of this work has been presented at Annual meeting of the Vision Science Society in May 2016^35^ and is documented in the PhD thesis of the first author ^24^. We thank Cheng Qiu and all the other members of the CPC laboratory and the Computational Neuroscience Initiative at the University of Pennsylvania for feedback and helpful discussions at various stages of the presented work.

## Supplementary Figures

### Individual subjects data for Experiment 1 and 2

Data of individual subjects are shown for Experiment 1 (Figs. S1 and S2) and Experiment 2 (Figs. S3 and S4). Figure S5 shows the estimated motor noise for each individual subject extracted from the motor training session (see Methods above).

**Figure S1:**
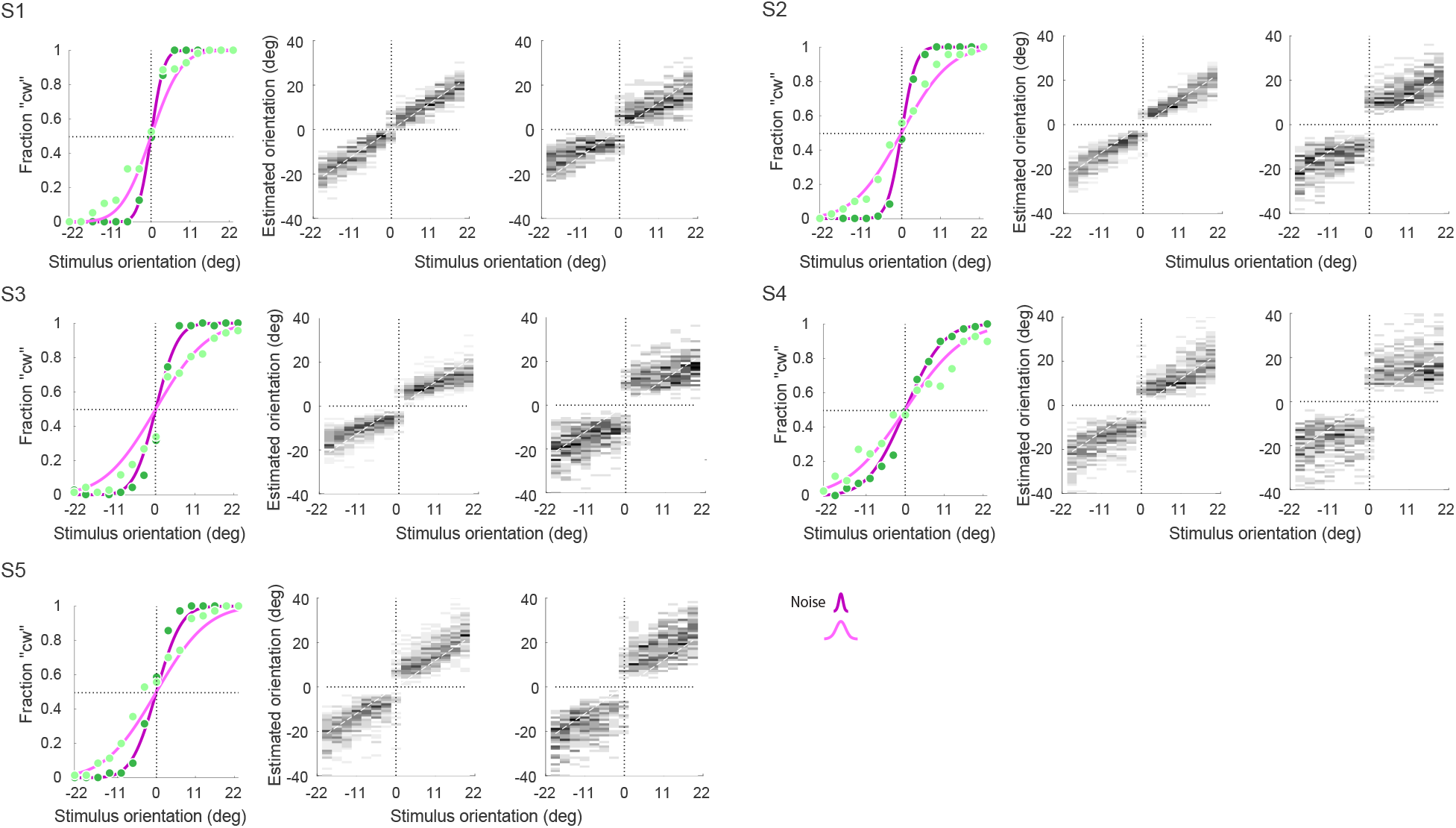
Experiment 1 data - individual subjects (correct trials).

**Figure S2:**
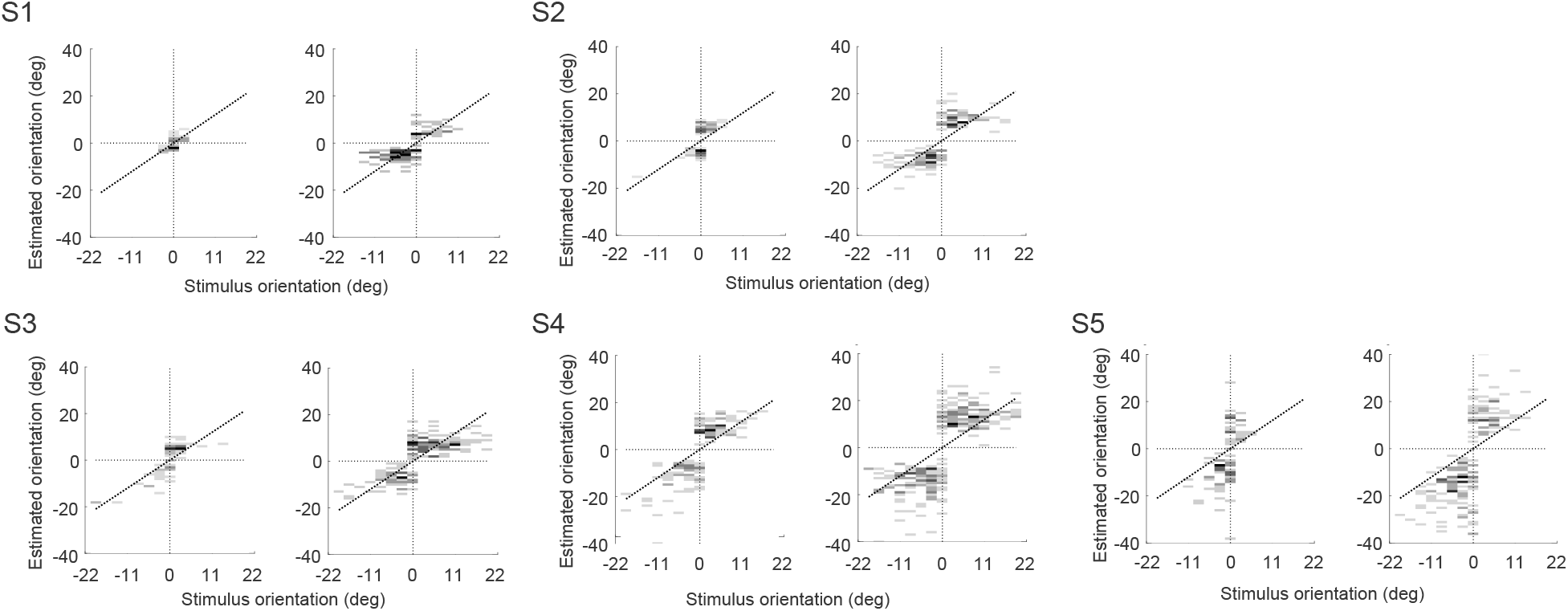
Experiment 1 data - individual subjects (incorrect trials).

**Figure S3:**
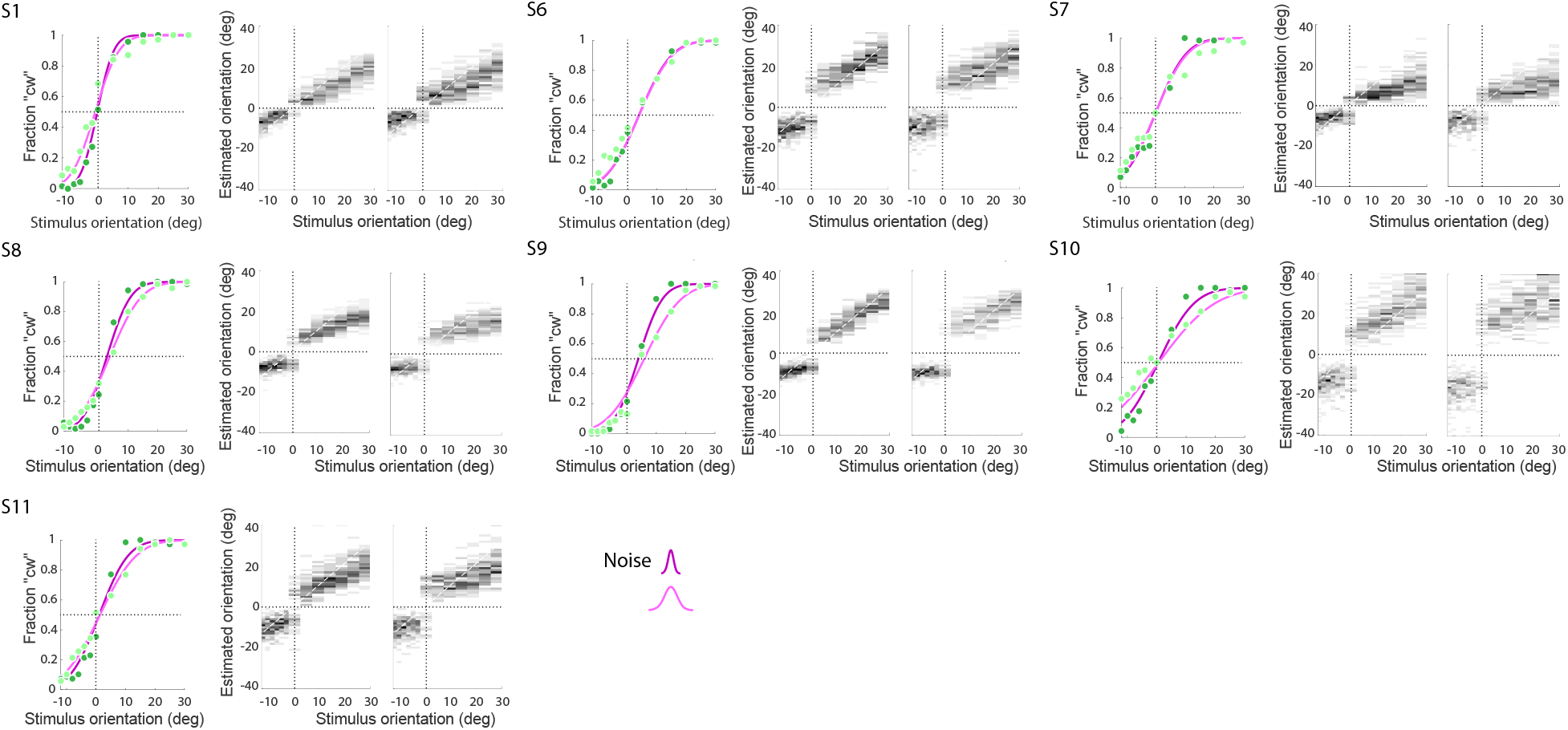
Experiment 2 data - individual subjects (correct trials).

**Figure S4:**
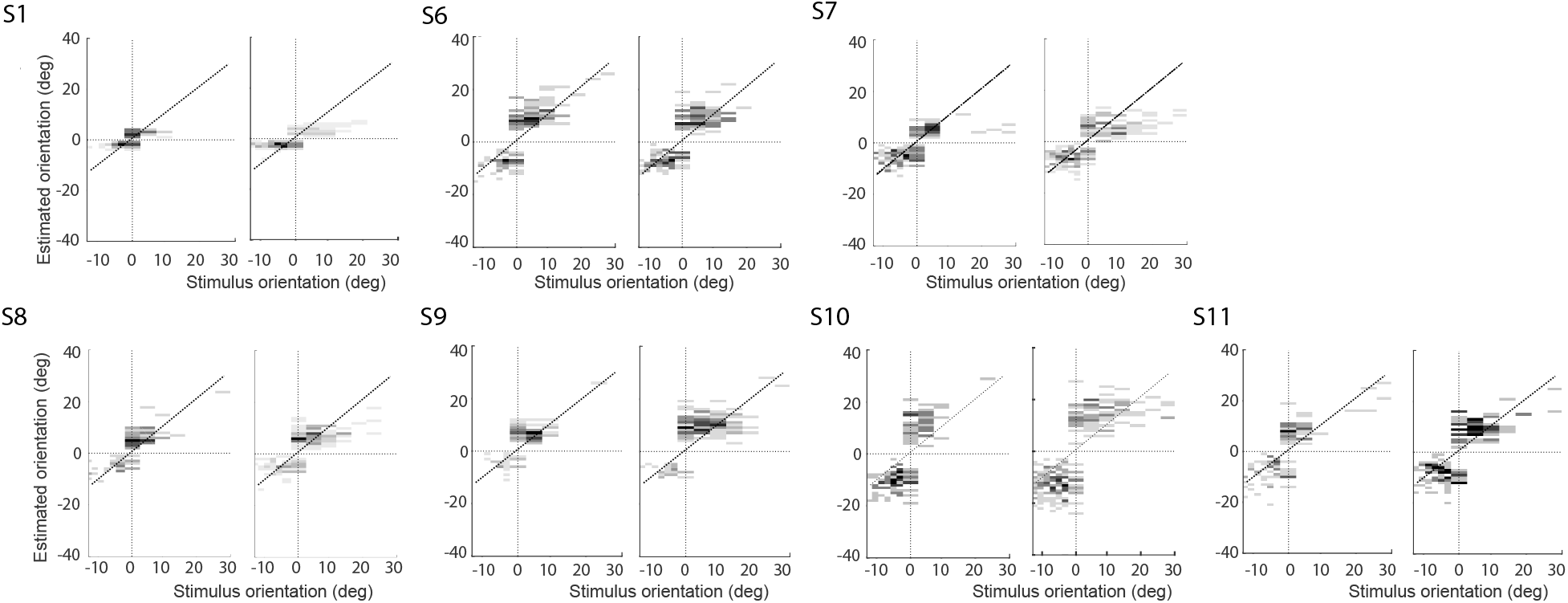
Experiment 2 data - individual subjects (incorrect trials).

**Figure S5:**
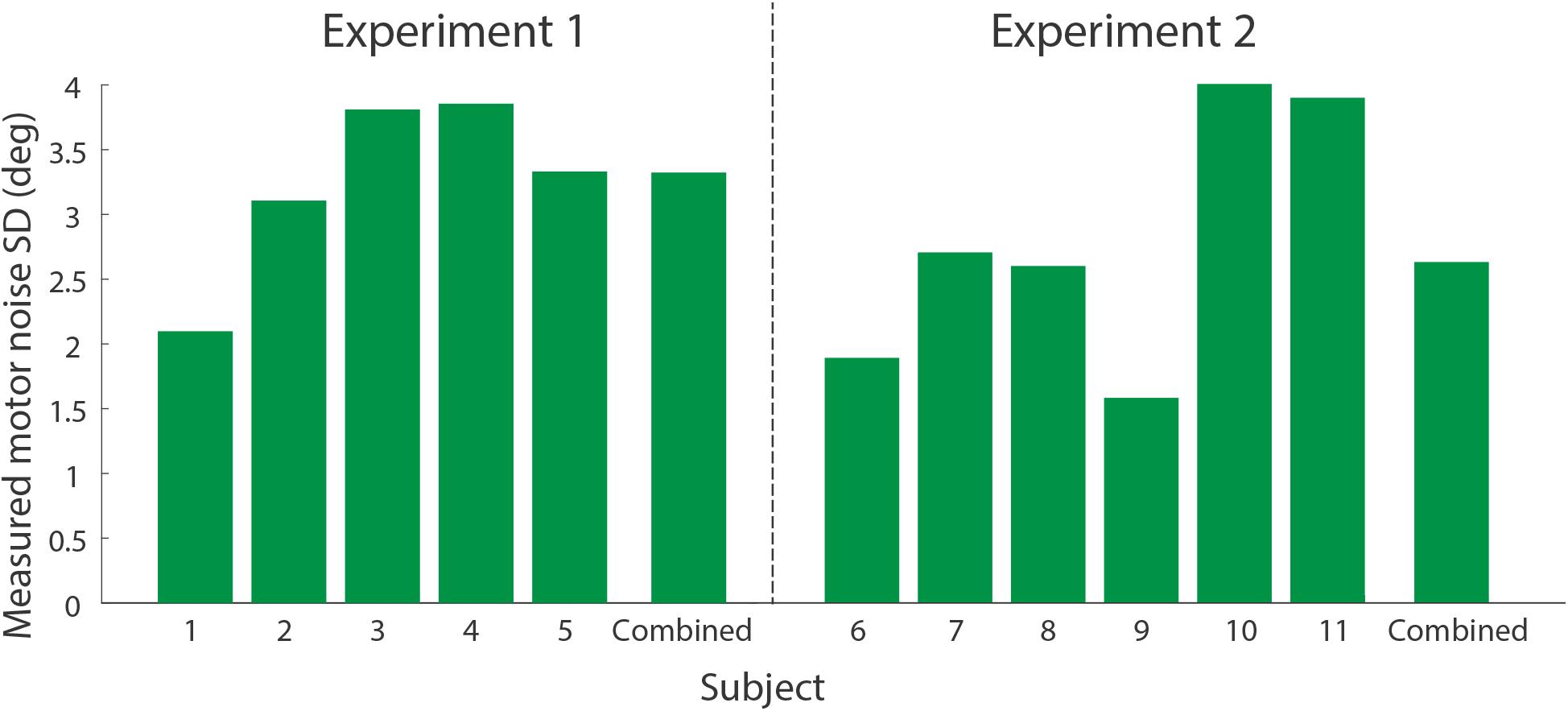
Motor noise of individual and combined subjects in Experiment 1 and 2.

### Model 2b (re-sampling) fit to correct trial data of individual subjects

Here we show fits and fit model parameters to correct trial data of individual subjects using the self-consistent observer model with re-sampling (Figs. S6 and S7). Figure S8 also shows the log-likelihood difference of the self-consistent observer model with and without conditioned resampling for each subject. For correct trials, fits of both versions of the model are essentially indistinguishable.

**Figure S6:**
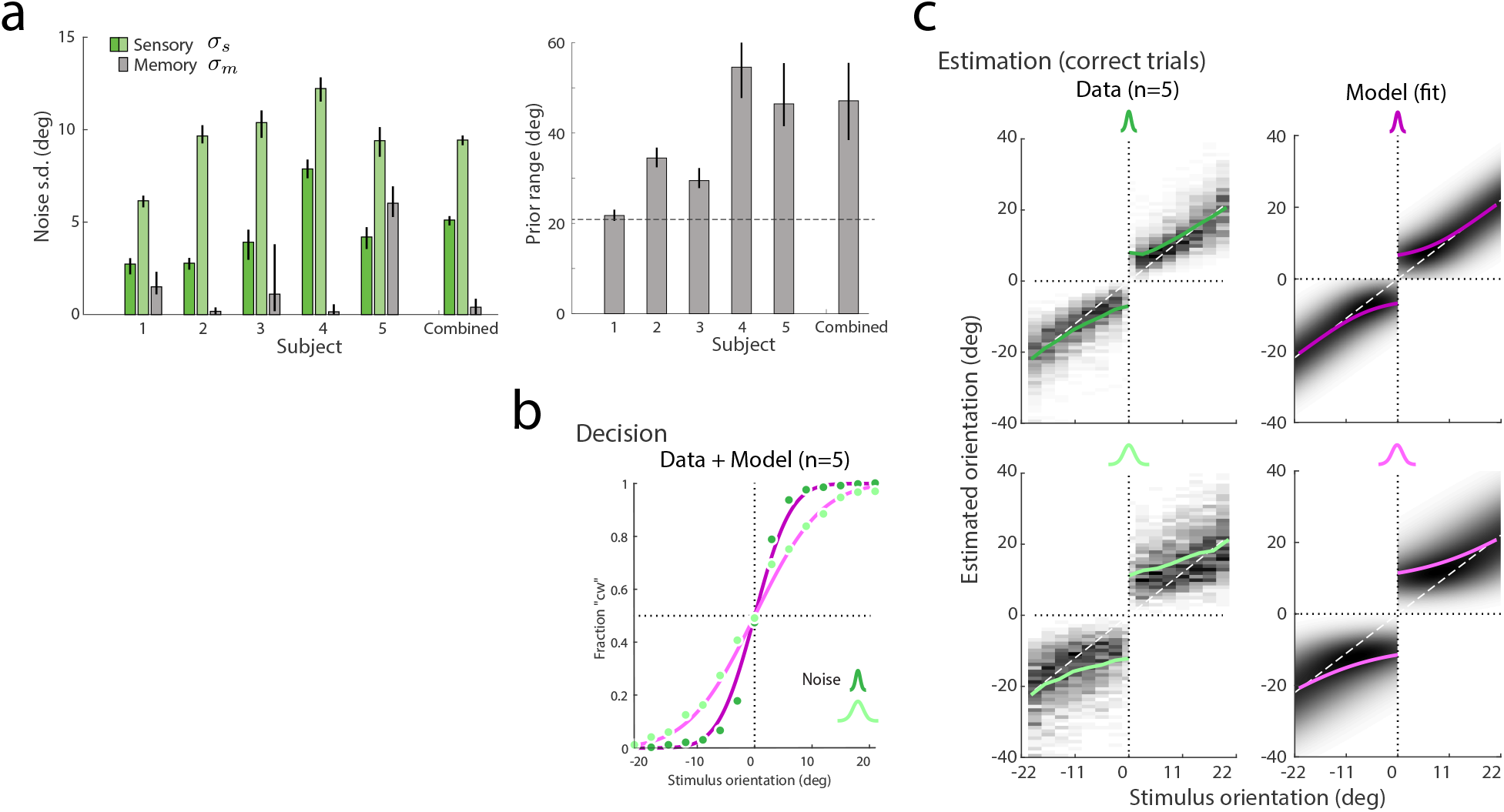
Model 2b fit to correct trials in Experiment 1.

**Figure S7:**
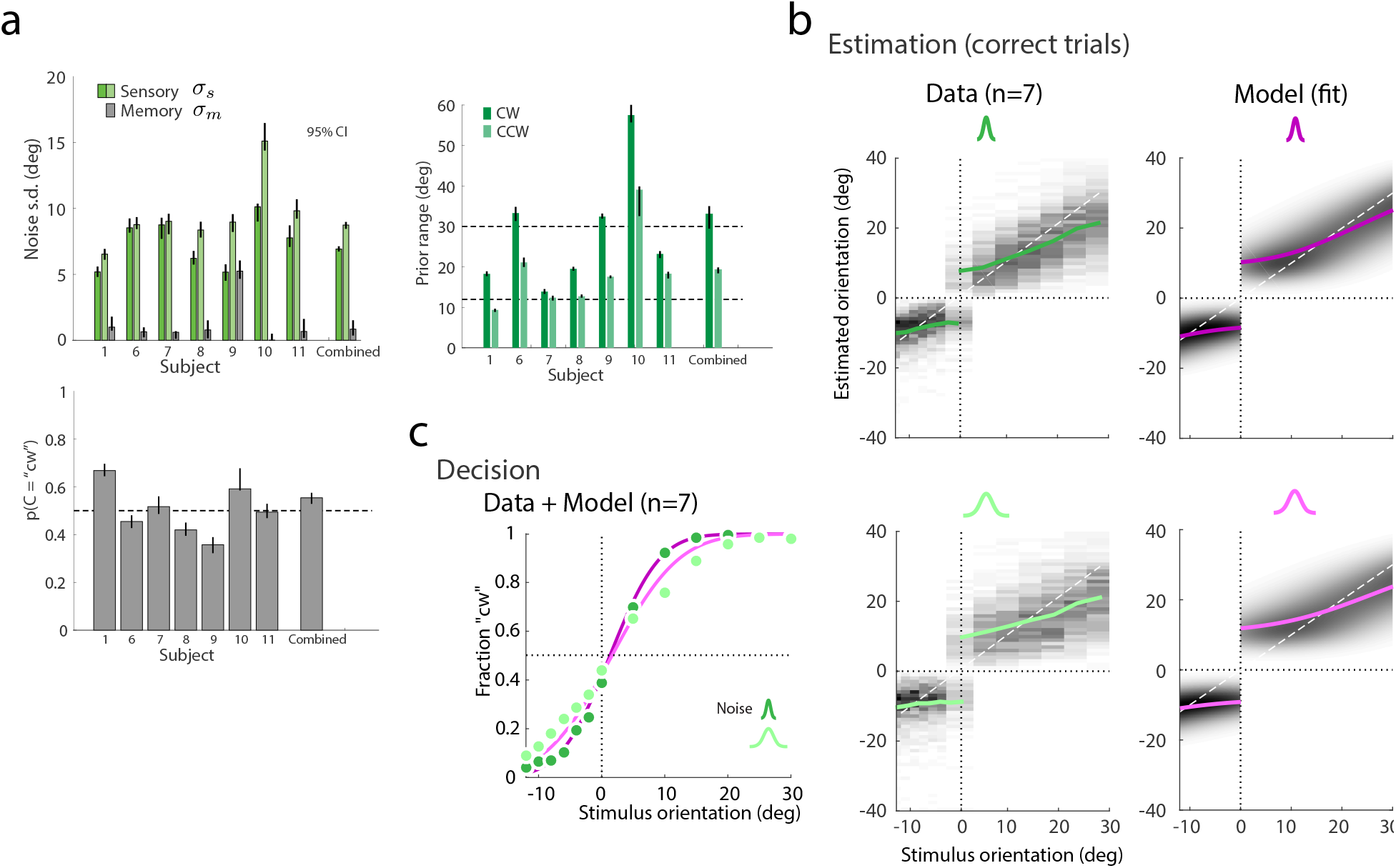
Model 2b fit to correct trials in Experiment 2.

**Figure S8:**
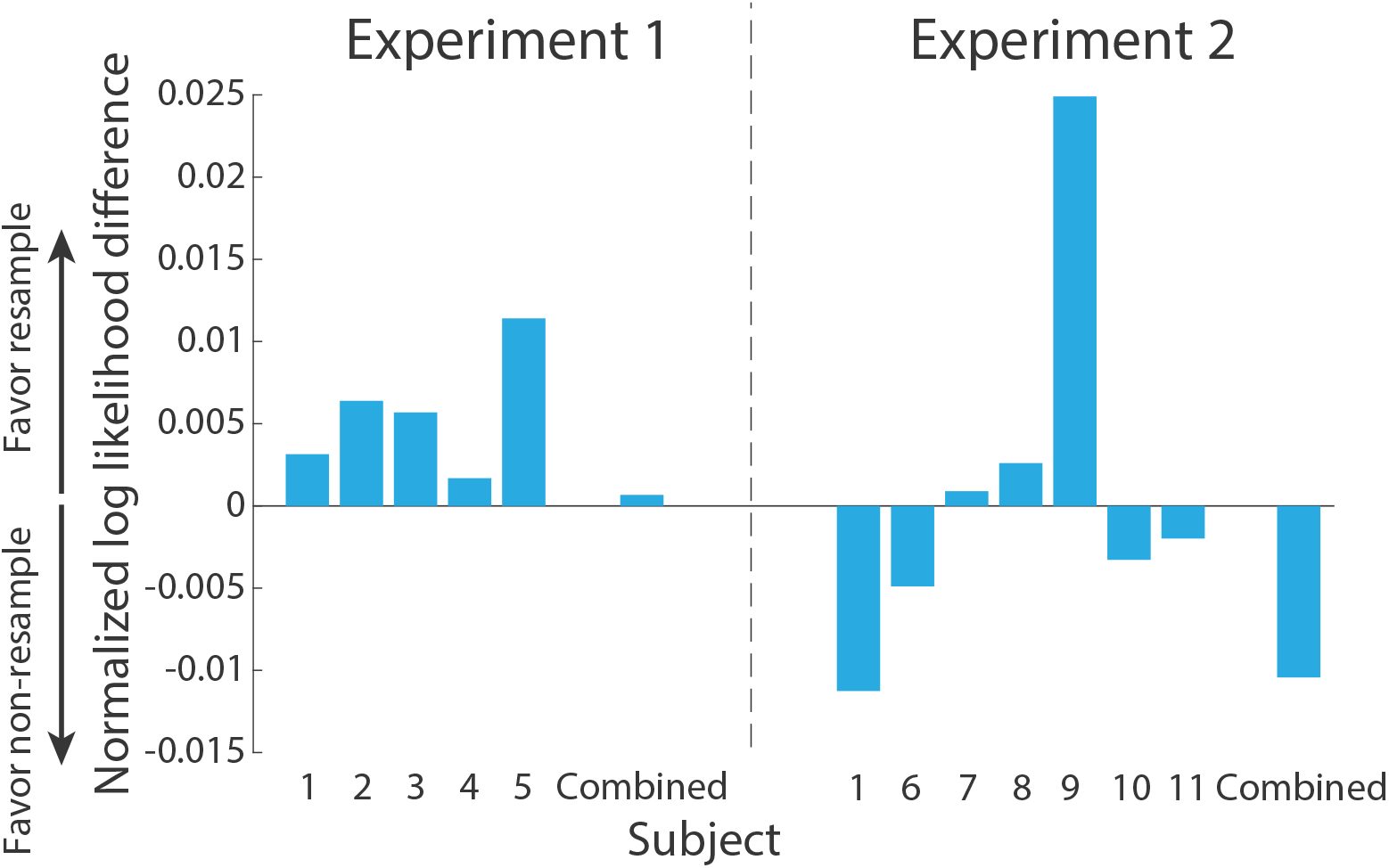
Normalized log likelihood difference between the resample and non-resample versions of the self-consistent observer model fit to correct trials in Experiment 1 and 2. The log likelihood values were normalized by the number of trials to make the comparison fair across subjects and experiments.

### Model 2b (re-sampling) fits to correct and predictions for incorrect trial data shown for each stimulus condition

**Figure S9:**
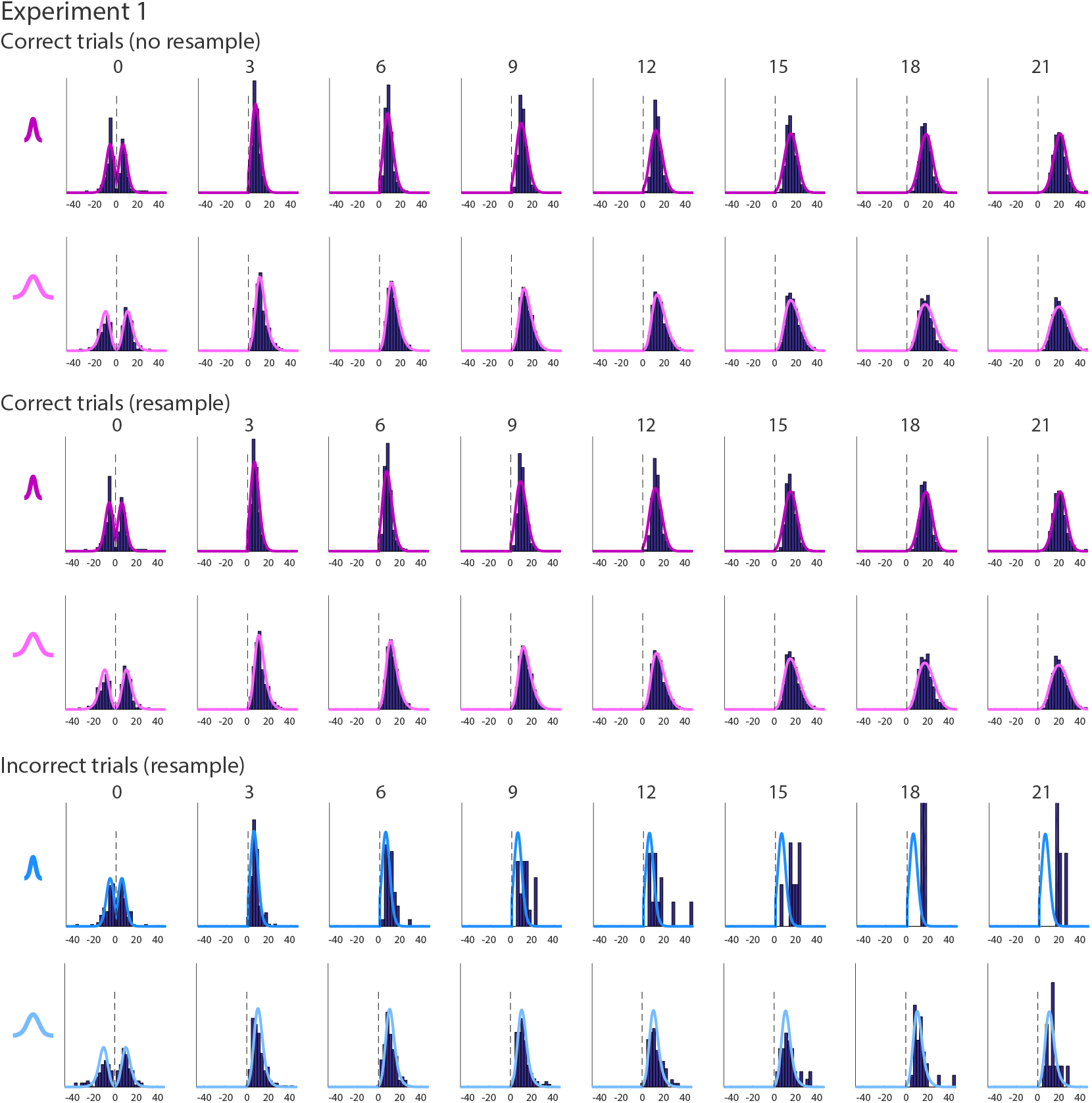
Model 2b prediction of incorrect trials in Experiment 1 (combined subject). For comparison, the top two rows show the fits of the self-consistent observer model without re-sampling. Note, histograms for stimulus orientations far away from the reference orientation only contain very few incorrect trials. Thus they do not provide a significant test of the model predictions.

**Figure S10:**
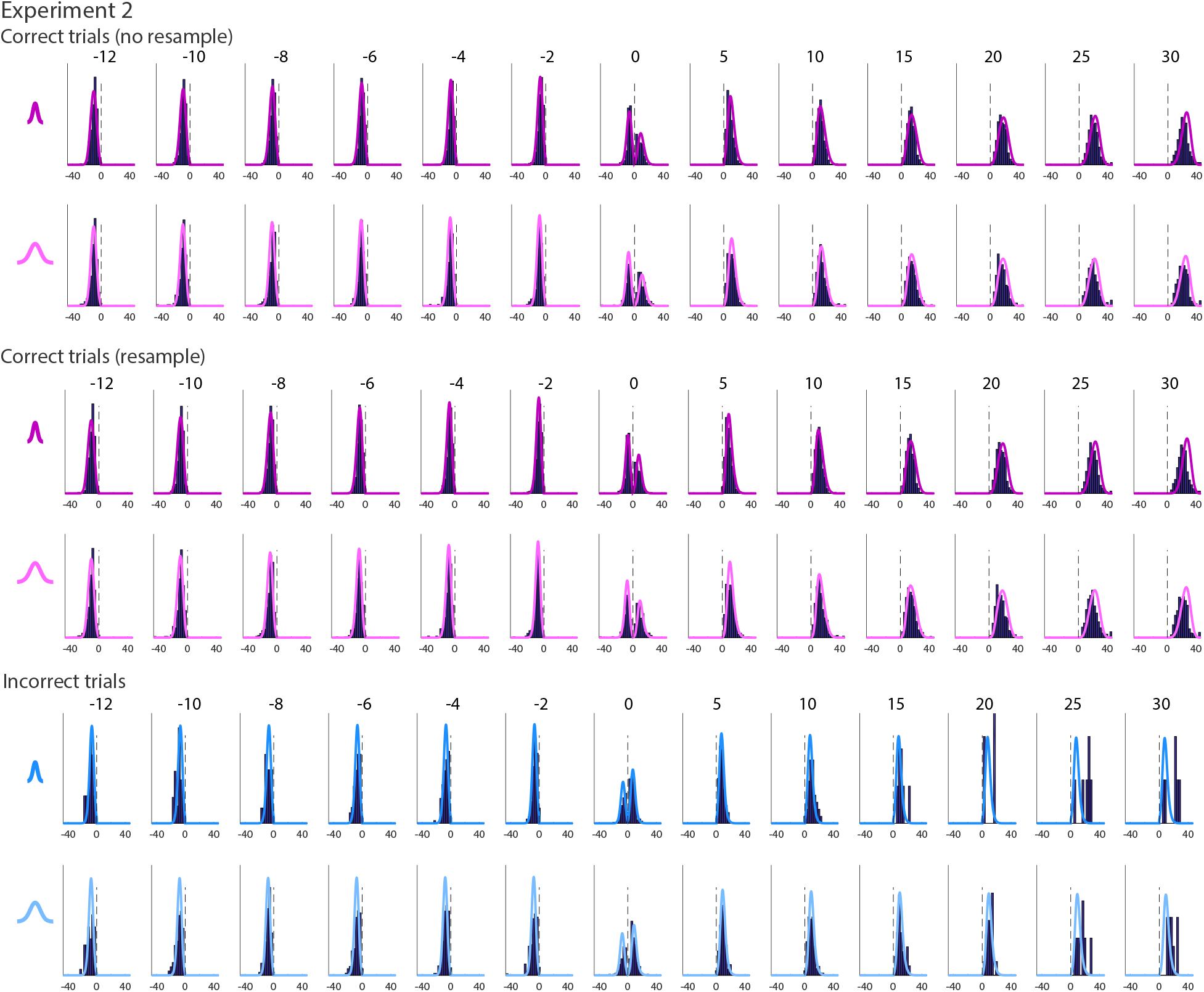
Model 2b prediction of incorrect trials in Experiment 2 (combined subject). For comparison, the top two rows show the fits of the self-consistent observer model without re-sampling. Note, histograms for stimulus orientations far away from the reference orientation only contain very few incorrect trials. Thus they do not provide a significant test of the model predictions.

## Notes

### Competing Interest Statement

The authors have declared no competing interest.

